# Antigenic characterization and pandemic risk assessment of North American H1 influenza A viruses circulating in swine

**DOI:** 10.1101/2022.05.04.490709

**Authors:** Divya Venkatesh, Tavis K. Anderson, J. Brian Kimble, Jennifer Chang, Sara Lopes, Carine K. Souza, Andrew Pekosz, Kathryn Shaw-Saliba, Richard E. Rothman, Kuan-Fu Chen, Nicola S. Lewis, Amy L. Vincent Baker

## Abstract

The first pandemic of the 21st century was caused by an H1N1 influenza A virus (IAV) introduced from pigs into humans, highlighting the importance of swine as reservoirs for pandemic viruses. Two major lineages of swine H1 circulate in North America: the 1A classical swine lineage (including the 2009 pandemic H1N1) and 1B human seasonal-like lineage. Here, we investigated the evolution of these H1 IAV lineages in North American swine and their potential pandemic risk. We assessed the antigenic distance between the HA of representative swine H1 and human seasonal vaccine strains (1978-2015) in hemagglutination inhibition (HI) assays using a panel of monovalent anti-sera raised in pigs. Antigenic cross-reactivity varied by strain but was associated with genetic distance. Generally, swine 1A lineage viruses that seeded the 2009 H1 pandemic were antigenically most similar to H1 pandemic vaccine strains, with the exception of viruses in the genetic clade 1A.1.1.3 that had a two-amino acid deletion mutation near the receptor-binding site, dramatically reducing antibody recognition. The swine 1B lineage strains, which arose from previously circulating (pre-2009 pandemic) human seasonal viruses, were more antigenically similar to pre-2009 human seasonal H1 vaccine viruses than post-2009 strains. Human population immunity was measured by cross-reactivity in HI assays to representative swine H1 strains. There was a broad range of titers against each swine strain that was not associated with age, sex, or location. However, there was almost no cross-reactivity in human sera to the 1A.1.1.3 and 1B.2.1 genetic clades of swine viruses, and the 1A.1.1.3 and 1B.2.1 clades were also the most antigenically distant from all human vaccine strains. Our data demonstrate that antigenic distances of representative swine strains from human vaccine strains represent a rational assessment of swine IAV for zoonotic risk research and pandemic preparedness prioritization.

**Importance:** Human H1 influenza A viruses (IAV) spread to pigs in North America, resulting in sustained circulation of two major groups of H1 viruses in swine. We quantified the genetic diversity of H1 in swine and measured antigenic phenotypes. We demonstrated that swine H1 lineages were significantly different from human vaccine strains and this antigenic dissimilarity increased over time as the viruses evolved in swine. Pandemic preparedness vaccine strains for human vaccines also demonstrated a loss in similarity with contemporary swine strains. Human sera revealed a range of responses to swine IAV, including two groups of viruses with little to no immunity. Surveillance and risk assessment of IAV diversity in pig populations are essential to detect strains with reduced immunity in humans, providing critical information for pandemic preparedness.

## Introduction

The first pandemic of the 21st century was caused by a strain of influenza A virus (IAV) that was introduced from pigs into humans, highlighting the importance of swine as reservoirs for pandemic influenza A viruses. Many strains of IAV that circulate in pigs are derived from repeated introductions of human IAV into swine populations (Ma *et al*., 2009; Nelson *et al*., 2012; Lewis *et al*., 2016; Rajao *et al*., 2018). Due to differences in population structure and movement, viruses in pigs may evolve differently from those circulating in humans (Lewis *et al*., 2016). Instead of strain replacement driven by targeted antibody responses to surface proteins and existing herd immunity, sub-populations of pigs with variable immunity and movement within production systems but limited movement of pigs between populations results in uneven antigenic change and multiple cocirculating lineages. In addition, reassortment with endemic swine IAV and further adaptation results in a diverse population of viruses (Gao *et al*., 2017; Rajao *et al*., 2019).

Three major lineages of viruses bearing the H1 hemagglutinin (HA) cocirculate in swine: these are named 1A, 1B, and 1C, linking the evolutionary history of the genes to common ancestral lineages (Anderson *et al*., 2016). H1-1A are derived from the “classical” 1918 human pandemic virus and have spread globally in swine. The 1A lineage also includes the H1 genetic clade that resulted in the 2009 H1N1 pandemic. Continued transmission of the pandemic virus in humans resulted in reintroduction of the virus into swine in multiple locations across the world, in some cases with onwards transmission within pigs (Nelson *et al*., 2015). In North America, genetic clades within the 1A lineage were previously classified into alpha (α), beta (β) and gamma (γ) and are now referred to as 1A.1.x, 1A.2, and 1A.3.x respectively (where x designates further subdivision). 1B viruses are derived from human seasonal H1 viruses that circulated prior to the 2009 pandemic and were introduced into pigs at different points in time. In the 1990s, 1B.1 viruses were first described in the United Kingdom (Brown *et al*., 1998). In the 2000s there were incursions of human seasonal H1 viruses into US swine (1B.2.1 (formerly δ2) and 1B.2.2 (formerly δ1) viruses) (Vincent *et al*., 2009; Lorusso *et al*., 2011), as well as introductions and onwards transmission in several other geographical locations including Chile, Argentina, Brazil, Australia, and Vietnam (Anderson *et al*., 2020). The 1C HA was derived from an avian H1N1 virus introduction which circulated in pigs in Europe in the 1970s, and then spread to Asia (Pensaert *et al*., 1981; Guan *et al*., 1996). A single detection of a 1C lineage HA gene was reported in Mexico (Mena *et al*., 2016), but no other 1C HA were identified in surveillance activities in North America.

Given relatively frequent transmission of IAV between humans and swine, and the ever-present risk of another swine-origin IAV pandemic, it is critical to objectively rank the zoonotic risk of the IAV circulating in pigs. Unlike avian influenza viruses, whose hemagglutinin (HA) commonly have receptor binding site (RBS) profiles for α2,3 linkage-type sialic acids prevalent in bird airway and intestinal cells, human-derived swine IAV HAs maintain the α2,6 linkage-type sialic acids dominant in human airways on re-introduction into pigs and consistently demonstrate the potential for replication and transmission in humans (Pulit-Penaloza *et al*., 2018; Sun *et al*., 2018; Kaplan *et al*., 2020). As the main target of host antibody responses, HAs that deviate significantly from those to which the human population have prior immunity, will pose a greater zoonotic risk if a variant virus was capable of human to human transmission. Further, it is plausible that cross-immunity and pandemic risk correlates with antigenic distance between a swine virus HA and human seasonal vaccine strains. The antigenic differences between swine and human HAs can be measured in the lab using binding assays such as the hemagglutination inhibition (HI) assay (Hirst, 1942). These data may subsequently be visualized and quantified using antigenic cartography to position viruses and sera in 2D or 3D “antigenic maps” such that map distances directly correspond to HI measurement (Lapedes and Farber, 2001; Smith *et al*., 2004). Consequently, characterizing the antigenic diversity of HA circulating in swine provides an objective assessment of the pandemic risk.

In this study, we used virus isolates, swine antisera and genetic sequences generated as part of the NIH CEIRS network pipeline to characterize currently circulating swine IAV in North America (USA, Mexico, and Canada). We measured antigenic distances between a representative sample of currently circulating H1 swine IAVs and H1 human vaccine strains since the 1970s. We also tested representative swine strains against pandemic preparedness vaccines, termed candidate vaccine viruses (CVV) using ferret sera. We then tested a subset of these viruses with age-stratified post-vaccination and post-exposure human sera to assess potential immunological cross-reactivity in the human population against diverse swine IAV strains. These data showed that antigenic distance from swine IAV to human vaccine strain is a rational measure to rank swine strains for pandemic risk, and should be linked with genomic epidemiology and existing risk assessment tools to inform public health pandemic preparedness measures.

## Materials and Methods

### Genetic analysis

All available swine H1 HA sequences from Canada, USA, and Mexico were downloaded from the Influenza Research Database (IRD) (Squires *et al*., 2012; Zhang *et al*., 2017) on 14 October 2019 (5517 sequences). To restrict the dataset to relevant field viruses, sequences of laboratory origin were excluded. All duplicate sequences and those with >30% ambiguous bases (“N”) were removed (4862 sequences). For visualization, sequences were down-sampled by nucleotide similarity using cd-hit (Cluster Database at High Identity with Tolerance) (Li and Godzik, 2006; Fu *et al*., 2012), removing sequences with > 98.0% sequence identity across the HA gene (375 sequences). To this dataset, the HA sequences of reference strains used in HI assays were added, while removing any duplicates to create the final dataset (432 sequences). Sequences were aligned using MAFFT v7.407 (Katoh *et al*., 2002; Katoh and Standley, 2013) and trimmed to starting ATG and ending stop codon. A maximum-likelihood (ML) phylogenetic tree was inferred using IQ-TREE v1.5.5 (Nguyen *et al*., 2015) following automatic model selection with statistical support determined using the Shimodaira-Hasegawa-like approximate Likelihood Ratio Test (aLRT, 1,000 replicates) (Guindon *et al*., 2010). Trees were plotted in R v3.6 using the ggtree package (Yu *et al*., 2017).

To characterize the genetic diversity, we selected 45 swine and human influenza A viruses (IAV) to test by HI against swine sera raised to 35 IAV strains, 26 against swine strains, 1 against an H1N1 variant strain, 1 against an early H1N1pdm09 human strain, and 7 against human vaccine strains. The test panel included some previously characterized swine H1N1 and H1N2 strains (Lorusso *et al*., 2011; Rajao *et al*., 2018). These antisera strains represented historical or contemporary clades of viruses from the US; for some international strains, we were restricted to viruses where an isolate was available. The 45 antigens were selected by classifying all H1 genes to genetic clade, generating an HA1 consensus sequence for each clade, and identifying the best matched field-strain to the consensus. In some cases, a genetic clade was represented more than once as it reflected a statistically supported monophyletic clade in the gene tree, or if detections of the clade came from different geographic locations. Strain names, subtypes, clades, and Genbank accession numbers can be found in **Supplementary Table S1**. No strains were selected from the 1A.2 (β) clade as it was infrequently detected in surveillance efforts with only 47 strains sequenced after 2015 (Arendsee *et al*., 2021). Similarly, we did not select strains from within the 1B.2.2.2 clade, despite representing a large number of HA genes in earlier years of the dataset, it was detected infrequently from 2016 to 2018, i.e., the numerical dominance changed from 1B.2.2.2 to 1B.2.2.1 in 2015 (Rajao *et al*., 2018; Zeller *et al*., 2018).

### Viruses and antisera production

Viruses were grown in Madin-Darby canine kidney cells with the exception of A/New Caledonia/20/1999 and A/Solomon Island/3/2006 which were grown in embryonated chicken eggs. Viruses used for serum production were clarified from cell culture supernatant, concentrated and UV inactivated. Each vaccine dose was approximately 128 HAU of antigen mixed 5:1 with oil-and-water adjuvant (Emulsigen D, MVP Laboratories, Inc. Ralston, NE).

Three-week-old cross-bred pigs free of IAV and IAV-antibodies were used for IAV antisera production. All pigs were treated with ceftiofur crystalline-free acid (Excede; Pfizer, New York, NY) and enrofloxacin (Baytril; Bayer Animal Health, Shawnee Mission, KS). Two pigs per virus received prepared vaccine via intramuscular injection. Two or three doses were given 2–3 weeks apart. When HI titers to homologous virus reached at least 1:160, pigs were humanely euthanized with pentobarbital (Fatal Plus, Vortech Pharmaceuticals, Dearborn, MI) for blood collection. Sera was obtained through centrifugation and stored at −□20□°C until use. Pigs were cared for in compliance with the Institutional Animal Care and Use Committee of the National Animal Disease Center, USDA-ARS.

Ferret antisera produced against CVV strains (1A.1.1 A/Ohio/24/2017, 1A.3.3.3 A/Ohio/09/2015, 1B.2.1 A/Ohio/35/2017, 1B.2.1 A/Michigan/383/2018, 1B.2.2.1 A/Iowa/32/2016,) and a contemporary human pdm lineage H1N1 seasonal vaccine strain (1A.3.3.2 A/Idaho/7/2018) were provided by the Virology, Surveillance and Diagnosis Branch, Influenza Division, Centers for Disease Control and Prevention (CDC), Atlanta, Georgia.

### Hemagglutination inhibition assays

Prior to HI testing, swine sera were heat inactivated at 56□°C for 30□min, then treated with a 20% suspension of kaolin (Sigma–Aldrich, St. Louis, MO) followed by adsorption with 0.5% turkey red blood cells (RBCs). HI assays were performed by testing the reference antisera panel against the panel of H1 viruses according to standard techniques (Kitikoon *et al*., 2014). Ferret antisera were treated and tested in a similar manner; however, 0.75% guinea pig RBCs were used for adsorption and in the HI assay. Human sera were similarly heat inactivated, then treated with RDE and adsorbed with 0.5% turkey RBCs. HI assays were performed according to standard techniques. Two biological replicates of each virus anti-sera were used. Geometric mean titers were obtained by log_2_ transformation of reciprocal titers divided by 10 and used for the analyses.

### Antigenic cartography

Cross-HI tables were mapped and merged in three dimensions using multi-dimensional scaling implemented at https://acmacs-web.antigenic-cartography.org/. Antigenic distances between viruses were calculated in antigenic units (AU), where 1 AU is equivalent to a 2-fold loss in HI cross-reactivity. As defined for the human seasonal vaccine strain update, we used 3 AU or ≥ 8-fold loss in cross-reactivity as a threshold of significant loss in cross-reactivity for the risk ranking system described below. Only antigen and serum points supported by four or more titer values, and antigens from strains isolated in 2012 or later were retained. Antigenic maps were exported from acmacs-web and antigenic distances between antigens generated in the 3D map were extracted from the maps and plotted using ggplot2 (Wickham, H, 2016:2) in R v3.6 (R Core Team, 2020). A subset of human seasonal vaccine strain antisera were used as replicates in each HI panel: the selected sera were determined by grouping by vaccine strains by decade, determining the distribution of antigens/sera in the map with cross-reactivity to each sera. The merged HI table with all antigens and sera used is in **Supplementary Table S2**.

### Prioritizing swine H1 strains for pandemic risk assessment

Given the large number of North American swine H1 IAV strains in our HI assays and limited volume of human and ferret sera samples, we applied a scoring system to objectively select swine strains for additional characterization against human sera. As our goal was to identify swine H1 strains that have higher pandemic potential, the ranking system determined whether a strain was representative of currently circulating H1 genetic diversity in swine, with an accompanying assessment of whether the strain was antigenically unique similar to a previous method for H3 strains (Souza *et al*., 2021). Our assessment scored the relative risk to humans of a swine H1 virus from the panel (query strain) as *Risk* = (*S*_*rep*_ + *S*_*match*_) + (*A*_*dist*_ + *A*_*ν*_), where the variables accounted for genetic sequence diversity, and the A variables accounted for antigenic diversity. *S*_*rep*_ was the number of BLASTp hits the query strain had with ≥97% amino acid to the dataset (BLASTp search of query strain against circulating strains) to identify the degree to which the tested strains represented all available swine H1 HA1;*S*_*match*_ was the number of BLASTp hits that the query strain had from the dataset BLASTp search of circulating strains against the query strain to indicate which of the test strains best represented the HA1 of all available swine H1; *A*_*dist*_ was the count of the number of times the query test strain was ≥ 3*AU* from human vaccine strains to estimate adult human immunity developed from seasonal H1 exposure or vaccination, and variance and outliers in the antigenic data controlled for by *A*_*ν*_ (standard deviation of all antigenic distance values, *σ* × 3). The dataset used for genetic sequence diversity included 2 years of U.S. H1 HA data and 10 years for non-U.S. H1 HA data, due to sparsity of recent data outside of the U.S. The highest scoring swine strains from each lineage and genetic clade were selected to be assayed against human sera (**Supplementary Table S3**).

### Human sera cohorts

Human convalescent sera representing different geographic locations and age were tested against contemporary swine IAV to assess human immunity. Two cohorts of human sera, post-infection with seasonal H1N1 and post-vaccination, were tested against the selected swine H1 viruses. The sera were collected in a multi-center cohort study in two hospitals in the US and three hospitals in Taiwan coordinated by the Johns Hopkins Center of Excellence in Influenza Research and Surveillance (Johns Hopkins University IRB00091667). In both cohorts, patient information was recorded by dedicated research coordinators following an informed consent process. The first cohort was composed of convalescent sera of influenza-infected individuals (n=10) from Taiwan and (n=10) from the US during the 2015-16 season collected at approximately 28 days post-infection with seasonal H1N1. Sera was collected from adult patients who presented at the hospital with influenza-like illness; this was defined as a documented or reported fever and any of the three respiratory symptoms (cough, headache, sore throat) within the past seven days of visiting the hospital. This cohort presented neutralizing antibody titers against the H1N1 vaccine strain (A/Michigan/45/2015). The second cohort was composed of sera collected from individuals (n=40) vaccinated in the fall of 2017 with a quadrivalent influenza vaccine at the Johns Hopkins Hospital employee occupational health clinic (Baltimore, MD, U.S.). These subjects presented neutralizing antibody titers against the H1N1 vaccine strain (A/Michigan/45/2015) with samples taken approximately 28 days post vaccination. Female and male were grouped by decade of age: 20-29, 30-39, 40-49, and >50.

Subsequently, a random number generator was used to select 5 per age group per gender. For HI results analysis, geometric mean HI titers were plotted against different birth cohorts (1946-76, 1977-88, 1989-96 based on marked changes in the antigenicity of circulating human H1 viruses – pre and post 1977 outbreak, and 1989 vaccine update) and Spearman’s rank correlation was calculated to assess the association between HI titers of human seasonal H1N1 vaccine strains and swine H1 strains. Figures were plotted using ggplot2 package in R v3.6. The raw titer dataset is provided in **Supplementary Table S4**.

### Data availability

Data associated with this study are available as supplemental material or posted at https://github.com/flu-crew/h1-risk-pipeline.

## Results

### Genetic analysis of North American swine H1 IAV strains

The maximum-likelihood phylogenetic tree inferred with representative HA gene sequences from human and swine hosts collected between 2012 and 2019 demonstrated concurrent circulation of two major lineages, 1A and 1B, in North America (Figure 1). Consistent with prior studies, the 1C or Eurasian-avian lineage was not detected among our sequences. The majority of HA data were collected in the contiguous USA through the USDA Influenza A Virus in Swine Surveillance System (**Figure S1)**.

**Figure 1.**
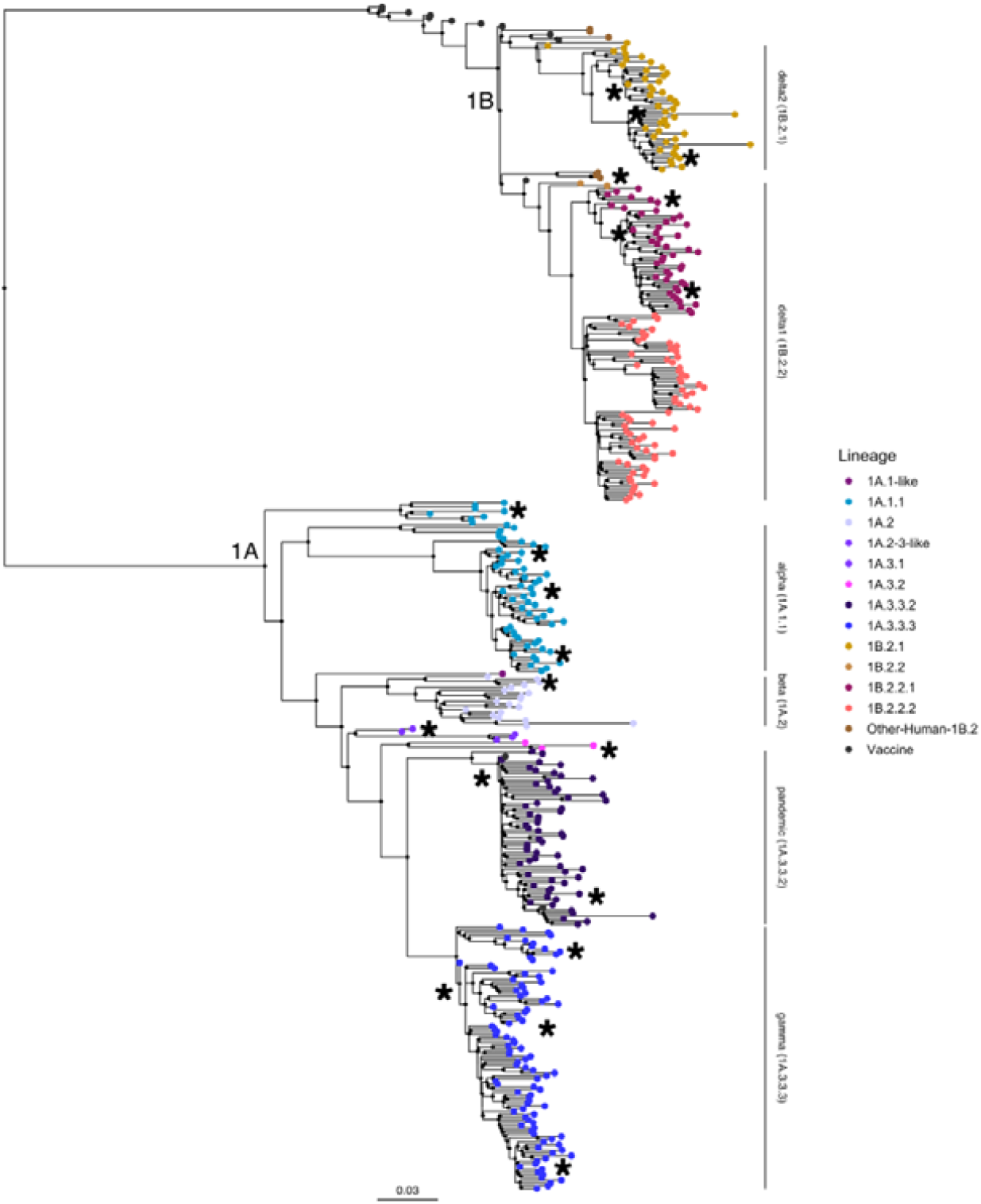
Phylogenetic tree of H1N1 influenza A viruses from North American swine populations during period of study (2012-2019). Swine IAV lineages are labelled on the right by annotated bars according to the global H1 clade classification. Monophyletic clades containing strains characterized in the risk pipeline are colored by clade at the tip point, swine strains selected for antigenic characterization are annotated by black asterisks, genetic clades with multiple representatives are only labelled once. All H1N1 human vaccine strain tip points, colored in black are included in the analysis but not labelled. The tree is mid-point rooted, all branch lengths are drawn to scale, and the scale bar indicates the number of nucleotide substitutions per site.

1A lineage viruses in USA, Mexico, and Canada circulated throughout the study period. The 1A.1.1 lineage viruses were detected within swine populations in Canada and USA, while the 1A.2 isolates were observed in Mexico and USA. The 1A.3.3.2 (H1N1pdm09 lineage) were detected in all three countries. The majority of 1A.3.3.2 transmission appeared to be derived from multiple introductions of human H1 into swine in each country rather than continued circulation within swine and transmission across countries. The 1A.3.3.3 lineage viruses were only detected in pig populations in the USA.

Swine 1B lineage viruses (1B.2.1 and 1B.2.2) were detected primarily in USA, although a few isolates were also detected in Mexico. The 1B.2.1 viruses detected in swine were not similar to any human vaccine strain in use in the early 2000s, a likely result of significant genetic evolution within pigs following the initial spillover. Genetically, the 1B.2.2.x clade viruses were most evolutionarily similar to a human seasonal strain, A/Michigan/2/2003 (approximate likelihood ratio test (aLRT) node support 99.7%). A/Solomon Islands/3/2006 and A/Brisbane/59/2007 formed a distinct sister clade to the 1B.2.1 strains and were the most similar, but low levels of public IAV sequence data in humans and swine prior to 2009 limit our ability to quantify genetic and antigenic relationships for this period.

### Antigenic evolution of swine H1 IAV in North America and distance to H1 human vaccine strains

Based on genetic representation and availability of virus isolates we selected a panel of swine H1 strains from each clade together with H1 human vaccine strains from 1977 to 2015. These strains were antigenically characterized in hemagglutinin inhibition (HI) assays against swine antisera, and antigenic maps were generated (**Figure 2**). Unlike previously-reported human or swine H3N2 antigenic analyses we did not detect antigenic clusters defined by a specific single or set of mutations or antigenic drift away from previously circulating strains in chronological order (Smith *et al*., 2004; Bedford *et al*., 2014; Abente *et al*., 2016; Lewis *et al*., 2016). Both 1A and 1B clades formed groups that were large, diffuse, and not chronologically oriented. However, there was evidence that antigenic distance among swine strains correlated with genetic distance (**Supplementary Figure S2)**. We also assessed potential risk of incursion into the human population, by measuring the distance from each swine strain to human vaccine strains (**Figure 3)**.

**Figure 2.**
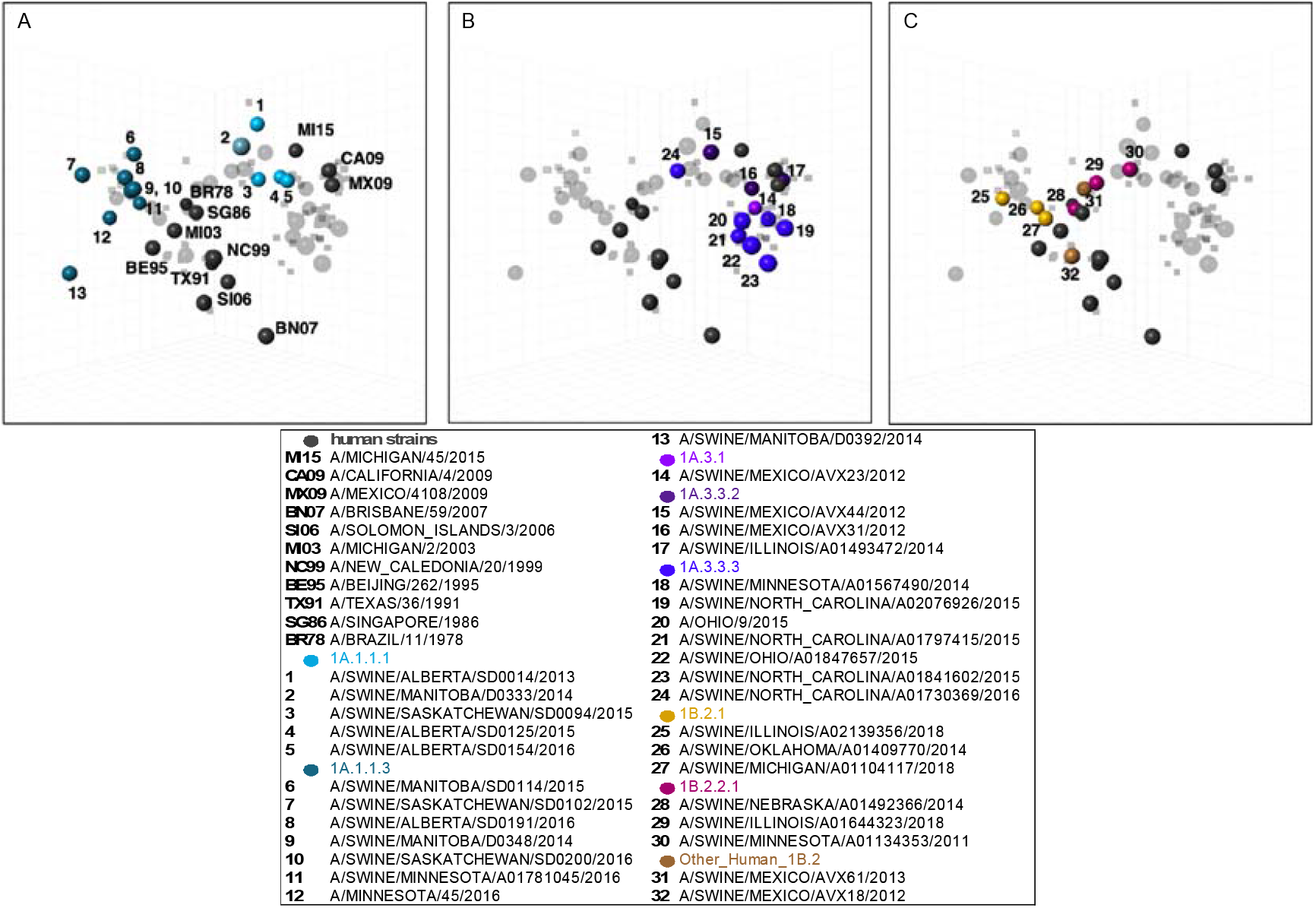
Antigenic relationships between human H1N1 vaccine strains and North American swine H1 strains. 3-dimensional maps are displayed in the same rotation in all panels. Human H1N1 vaccine strains (grey) are shown in all panels but labelled only in panel A with abbreviated strain names. North American swine H1N1 strains are split by lineages (A) 1A.1.1.x, (B) 1A.3.3.x and (C) 1B.2.x. Each sphere in the maps represents a strain colored according to the phylogenetic clade as described in the legend. All swine strains displayed significant antigenic heterogeneity among and between lineages. Human H1N1pdm09 strains (CA09, MX09, MI15) are antigenically distant from previous human vaccine strains and closer to the swine 1A strains (A, B). Antigens 8-13 in panel A with a double deletion in addition to other amino acid changes via drift, which likely explains their distance from the rest of the 1A strains. The antigenic variability of 1B.2.x (C) is consistent with multiple spillovers from contemporary pre-pandemic human seasonal strains (BE95-MI03).

**Figure 3.**
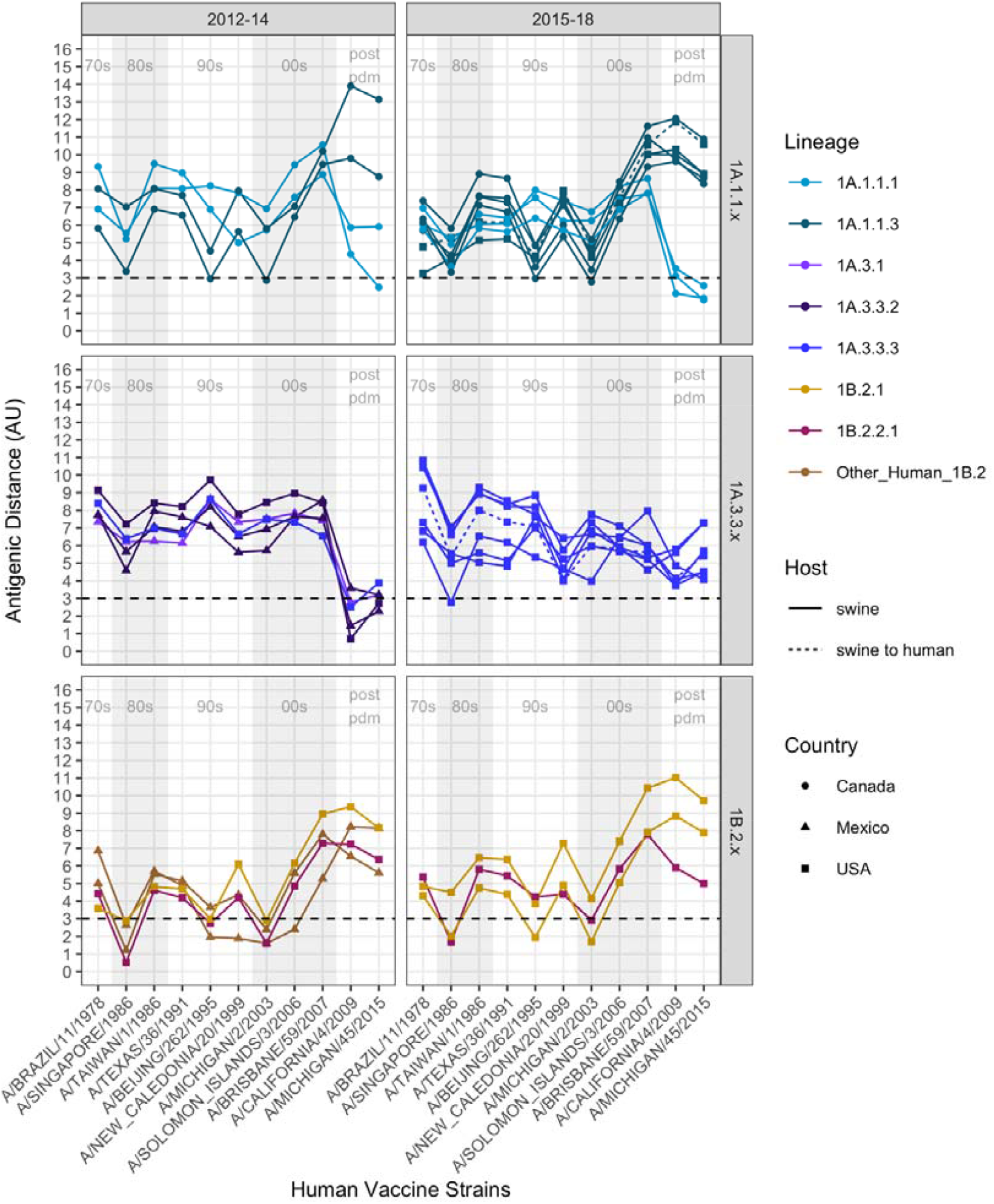
Antigenic distances between contemporary swine H1N1 strains and human seasonal H1N1 vaccine strains. Graphs are divided by swine strain lineage (1A.1.1.x, 1A.3.3.x and 1B.2.x) and were grouped using year of isolation to facilitate visualization. Column shades in light grey and white indicate human H1N1 vaccine strains by decade as listed along the x-axis of the lower graphs in chronological order (1978-2015). Antigenic distance between the swine strain and human vaccine strain are plotted on the Y-axis. Strains are colored by phylogenetic lineage as in Figure 1 and 2. The shape of the strain represents its country of origin (Canada, circle; Mexico, triangle; USA, square). The dotted line indicates significant antigenic distance (3 AU, ∼8-fold loss in HI titer) between the swine H1 strains on the y-axis and human seasonal H1N1 vaccine strains along the x-axis. In general, 1A lineages circulating in swine since the 1918 pandemic had variable distance to vaccine strains prior to 2009, but were closer to the H1N1pdm09 vaccine strains. The double deletion lineages in 1A.1.1.3 deviated from 1A.1.1.1 and were antigenically highly distant from most human vaccine strains, especially H1N1pdm09 strains. 1B lineages conversely are closer to pre-pandemic vaccine strains and >3AU from post-pandemic vaccine strains.

Antigenically, the 1A.1.x and the 1A.3.x lineage viruses in pigs were distinct from the 1B lineage strains (Figure 2 A-C). The 1A.1.1.1 lineage viruses circulating in Canadian pigs were more antigenically similar (1.75 – 5.92 AU) to the 2009-pdm like vaccines than the 1A.1.1.3 lineage viruses (8.34 – 11.86 AU) (Figure 2A and 3). The positioning of 1A.1.1.3 lineage viruses with deletions at positions 129 and 130 were antigenically closer to the swine 1B and related human vaccine strains in the map; the 1B lineage strains have a single deletion at residue 130 of mature H1 peptide. The 1A.3.3.2 lineage swine viruses and the 2009 pandemic and subsequent seasonal human vaccine strains were also antigenically distant from the seasonal influenza strains circulating in humans prior to 2009 (6.19 – 9.62 AU) and 1B lineage swine strains seeded by the pre-2009 human strains. Unlike other genetic clades within the 1A lineage, 1A.3.x viruses were antigenically more similar to the human H1N1pdm09 viruses, but with significant strain to strain variation (0.71 – 7.29 AU, **Figure 2B and 3**). The antigenic map point positions reflected antigenic drift that occurred in humans that resulted in the updated 2015 H1N1 pandemic vaccine strain (AU 2.29 distance to previous vaccine A/California/4/2009). In swine, however, our selected viruses were detected between 2009-14, and all of these HA genes were antigenically similar to the 2009 human vaccine strains. The other dominant circulating clade within 1A.3 were the 1A.3.3.3 strains that are geographically restricted to USA and showed significant genetic variation and corresponding antigenic diversity (range 3 to 11 AU from the tested human vaccine strains) likely driven by changes in the putative antigenic sites of H1 (Caton *et al*., 1982) (**Supplementary Figure S3A-D)**.

There was antigenic distinction between the 1B.2.2.x and 1B.2.1 viruses in the antigenic map, consistent with the phylogeny and accounting for the separate introductions of human seasonal influenza viruses into pigs in the early 2000s (**Figure 2C**). The antigenic distances of 1B strains against all human vaccine strains (**Figure 3**) showed that the most antigenically similar human vaccine strains were A/Beijing/262/1995 and A/Singapore/1986, but the closest antigenic distance was to the seasonal field strain, A/Michigan/2/2003 (1.6-4.4 AU). While both 1B.2.2.x and 1B.2.1 strains showed increased antigenic distance to the H1N1pdm09 human strains, the distance was greater in 1B.2.1 relative to the 1B.2.2.x strains.

### Candidate vaccine virus reactivity to contemporary swine H1 strains

To assess the potential of CVV and human seasonal H1 vaccines to protect against contemporary clades in pigs we tested representative swine strains of each of the regularly detected H1 clades from the 1A and 1B lineages against reference ferret antisera. Sera were raised against a human seasonal H1N1 vaccine strain A/Idaho/7/2018 and CVVs A/Ohio/24/2017 (1A.1.1.3), A/Ohio/9/2015 (1A.3.3.3), A/Iowa/32/2016 (1B.2.2.1), A/Ohio/35/2017 and A/Michigan/383/2018 (1B.2.1) (Table 2A and B). Swine viruses from clades 1A.1.1.x (including those that have the two amino acid deletion), 1A.3.3.3, and 1A.3.3.2 (H1N1pdm09 HA clade) showed moderate to significant cross-reactivity with sera raised against their respective within-clade CVV/human vaccine strains with ranges of 3-5-fold decreases in the HI assay (Table 2A). The 1A.2 and 1A.2.3-like viruses do not have a within-clade CVV or vaccine strain; and they demonstrated limited antigenic similarity to other 1A CVVs, but retained cross-reactivity with the high-titer A/Idaho/7/2018 vaccine strain. 1B.2.1 and 1B.2.2.1 viruses retained some cross-reactivity to sera from lineage-specific CVVs, even if at 2-3-fold decrease (Table 2B). 1B.2.2.2 viruses with no associated within-clade CVV reacted poorly with other 1B CVVs and had barely detectable titers but represent a minor clade detected in surveillance in the US.

**Table 1A and B.** Ferret anti-sera against human seasonal vaccine strains or candidate virus vaccines (CVVs) demonstrated variable recognition of swine H1. Hemagglutination inhibition (HI) titers of CVV or human seasonal vaccine anti-sera against contemporary North American swine H1 1A.1.x (A) and 1B.2.x (B) lineages. Homologous titers are highlighted in grey, and boxes mark titers against swine viruses of the same lineage.

|  |  |  | A/Ohio/24/2017 | IDCDC-RG59 A/Ohio/24/2017-like | A/Ohio/9/2015 RG48A | A/Idaho/7/2018 |
| --- | --- | --- | --- | --- | --- | --- |
| IDCDC-RG59 A/Ohio/24/2017-like | CVV | 1A.1.1 | 5120 | 2560 | <10 | 1280 |
| A/swine/Iowa/A02478635/2019 | Test | 1A.1.1 | 160 | 160 | 10 | 80 |
| A/swine/Oklahoma/A02245237/2019 | Test | 1A.2 | 40 | 20 | 320 | 2560 |
| A/swine/Texas/A01104132/2019 | Test | 1A.2-3-like | 20 | 20 | 80 | 1280 |
| A/Ohio/9/2015 | CVV | 1A.3.3.3 | <10 | <10 | 1280 | 10 |
| A/swine/North Carolina/A02478571/2019 | Test | 1A.3.3.3 | <10 | 10 | 1280 | 20 |
| A/swine/Iowa/A01731653/2016 | Test | 1A.3.3.3 | 10 | <10 | 160 | 20 |
| A/swine/Nebraska/A02214231/2017 | Test | 1A.3.3.3 | 10 | <10 | 320 | 10 |
| A/Idaho/7/2018 | Human vaccine | 1A.3.3.2 | 80 | 160 | 160 | 10240 |
| A/swine/Utah/A02432386/2019 | Test | 1A.3.3.2 | 80 | 40 | 160 | 5120 |

B
|  |  |  | A/Iowa/32/2016 | A/Ohio/35/2017 | A/Michigan/383/2018 |
| --- | --- | --- | --- | --- | --- |
| A/Iowa/32/2016 | CVV | 1B.2.2.1 | 640 | 20 | 20 |
| A/swine/South Dakota/A01481702/2014 | Test | 1B.2.2.1 | 80 | 20 | 80 |
| A/swine/Minnesota/A02478597/2019 | Test | 1B.2.2.2 | 40 | 20 | 20 |
| A/Ohio/35/2017 | CVV | 1B.2.1 | 80 | 640 | 160 |
| A/Michigan/383/2018 | CVV | 1B.2.1 | 40 | 160 | 1280 |
| A/swine/Illinois/A02139356/2018 | Test | 1B.2.1 | 20 | 320 | 1280 |

There were zoonotic events in humans from the 1A.1.1.3 clade, in 2016 (A/Minnesota/45/2016) and 2017 (A/Ohio/24/2017), with a CVV selected based on the 2017 strain. Sera raised in ferrets against the 2017 A/Ohio/24/2017 CVV reacted to the 2019 isolate from this clade in HI assay (**Table 2A**), but with a >4-fold decrease in cross-reactivity. The reduction in reactivity may be associated with two amino acid substitutions, one in a putative antigenic site Sb (H1-G153D) and a second in the putative receptor binding site (H1-T222A). Viruses from lineages such as the 1A.3.3.3 resulted in human zoonotic cases, with one selected as a CVV, A/Ohio/09/2015. Sera raised against this virus appears to retain cross-reactivity to contemporary viruses from the same 1A.3.3.3 clade (A/swine/North Carolina/A02478571/2019) but titers were decreased by 2-3 fold for other viruses within the 1A.3.3.3 clade.

### Human population immunity against swine H1 IAV

To understand the implications of IAV genetic and antigenic diversity in swine for human population immunity, we tested a subset of human vaccine and swine viruses against sera from age-stratified human cohorts. One strain from each lineage was chosen according to the risk-ranking scoring system described in the methods section (see **Supplementary Table S3**). One set of sera was collected from people vaccinated against the pandemic lineage virus A/Michigan/45/2015 (n=40, age: 22-68) at the Johns Hopkins Hospital, Baltimore, MD, US (post-vaccination cohort); and a second set of sera was collected from patients with confirmed influenza (subtype unknown: n=20, age: 21-71) in Taiwan (n=10) or in Baltimore (n=10) (post-exposure cohort). The cohorts included adults of both sexes between the ages of 21 and 71. The sera were tested against human vaccine strains: A/Beijing/262/1995 (BE95) and A/Michigan/45/2015 (MI15); and the following strains A/swine/Nebraska/A01492366/2014 (NE14 1B.2.2.1), A/swine/Illinois/A02139356/2018 (IL18 1B.2.1), A/swine/Mexico/AVX18/2012 (MX12 OtHu1B.2), A/Swine/Saskatchewan/SD0094/2015 (SK15 1A.1.1.1), A/Swine/Minnesota/A01781045/2016 (MN16 1A.1.1.3),A/swine/Iowa/A01731653/2016 (IA16 1A.3.3.3), and A/Swine/Mexico/AVX23/2012 (MX12 1A.3.1).

We did not detect major differences between titers from the post-exposure (Figure 4A) and post-vaccination (Figure 4A) cohorts in overall responses to different strains (**Supplementary Table S4**). There were heterogenous responses to most viruses, with a broad range of titers against each strain. The MN16 1A.1.1.3 and IL18 1B.2.1 strains were exceptions; as there appeared to be no cross-reactivity in almost all subjects (GMT less than 2: and 4B). These data mirror the swine sera HI data (Figure 3), where the MN16 1A.1.1.3 strain was antigenically distant to both pandemic and pre-pandemic human seasonal strains circulating in the 2000s with an average distance AU 9.6 and 7.07 respectively; and was also significantly drifted from human strains from prior decades (average distance AU 4.17 and 5.57 for vaccines used in the 1970-1980s and 1990s respectively). The IL18 1B.2.1 strain had a similar trend: with the most similar human vaccine strain A/Beijing/262/1995 at 3.85 AU distance, and above the cut-off used when human vaccine strains are recommended for revision due to decreases in cross-reactivity to antisera (**Supplementary Table S3**). We parsed the HI data by age to test whether the range of cross-reactive titers could be separated into high or low reactors (**Figure 5**). The data showed that birth years split into categories of 1946 to 1976, 1977 to 1978, and 1989 to 1996 were not associated with titer response to swine or human IAV strain. Regardless of the birth periods, there were both high and low reactors to each of the viruses with the exception of MN16 1A.1.1.3 which had consistently low reactions (Figure 5, Figure S4). The IL18 1B.2.1 demonstrated a unique pattern, with two individual samples in the 1977-1988 birth year cohort exhibiting a high response to the virus (Figure 5, Figure S4B).

**Figure 4.**
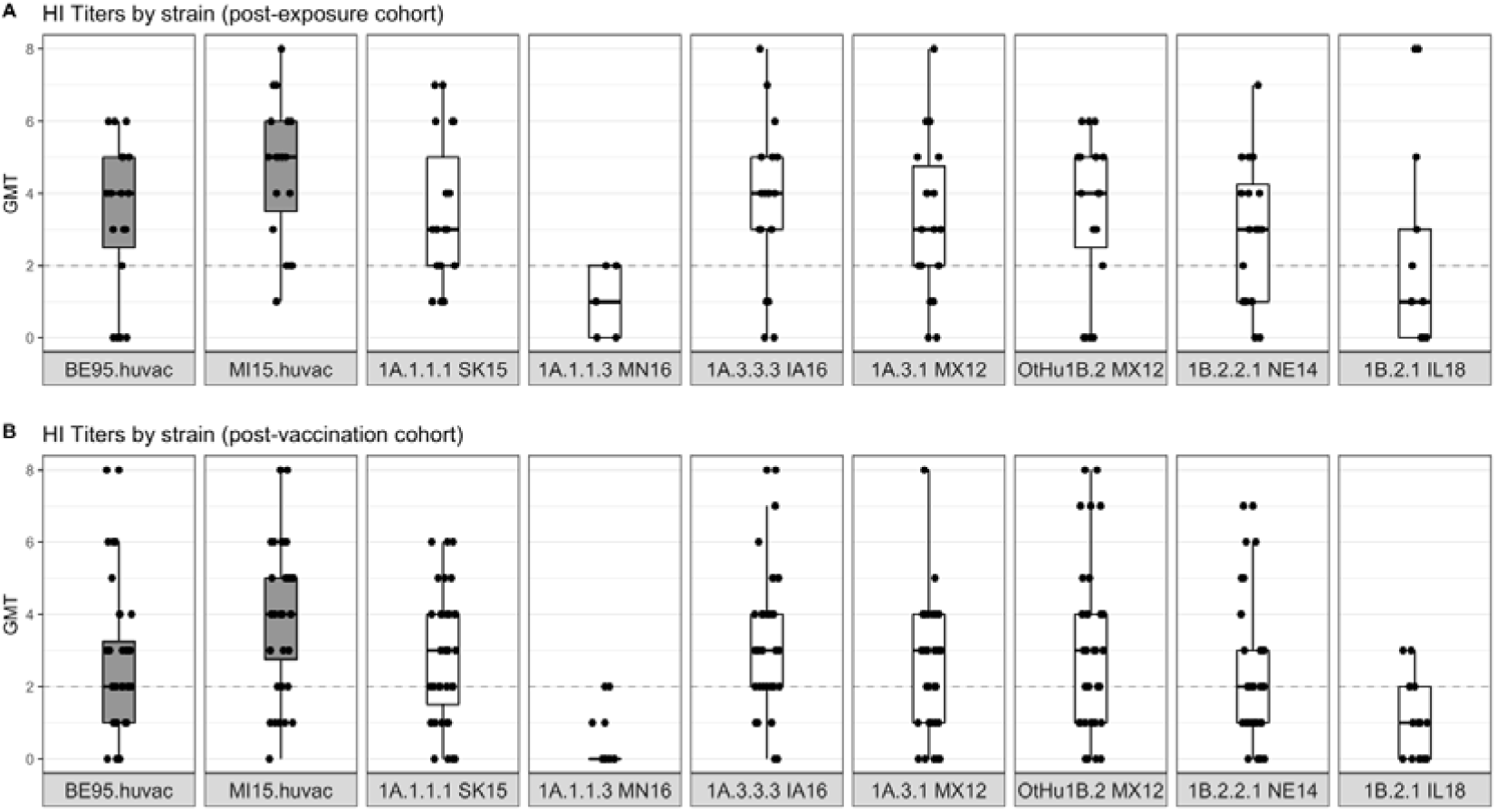
Human convalescent (A) and post-vaccination (B) sera geometric mean log _2_ hemagglutination inhibition (HI) titers against North American H1N1 swine strains. Box plots show the median of aggregated HI titers against H1N1 strains with 5th and 95th percentile and standard deviation. Each dot represents the GMT: geometric mean titer, log_2_ (HI titer /10) of human sera on the y-axis against each strain (shown on x-axis). Boxplots in grey indicate the human H1N1 vaccine strains and boxplots in white indicate swine H1 strains. The grey dotted line indicates the minimum positive HI titer threshold (≥40 or 2). Human sera show a range of titers against most swine and human vaccine strains, with the exception of MN16 (1A.1.1.3) and IL18 (1B.2.1) strains to which there was little to no cross-reactivity.

**Figure 5.**
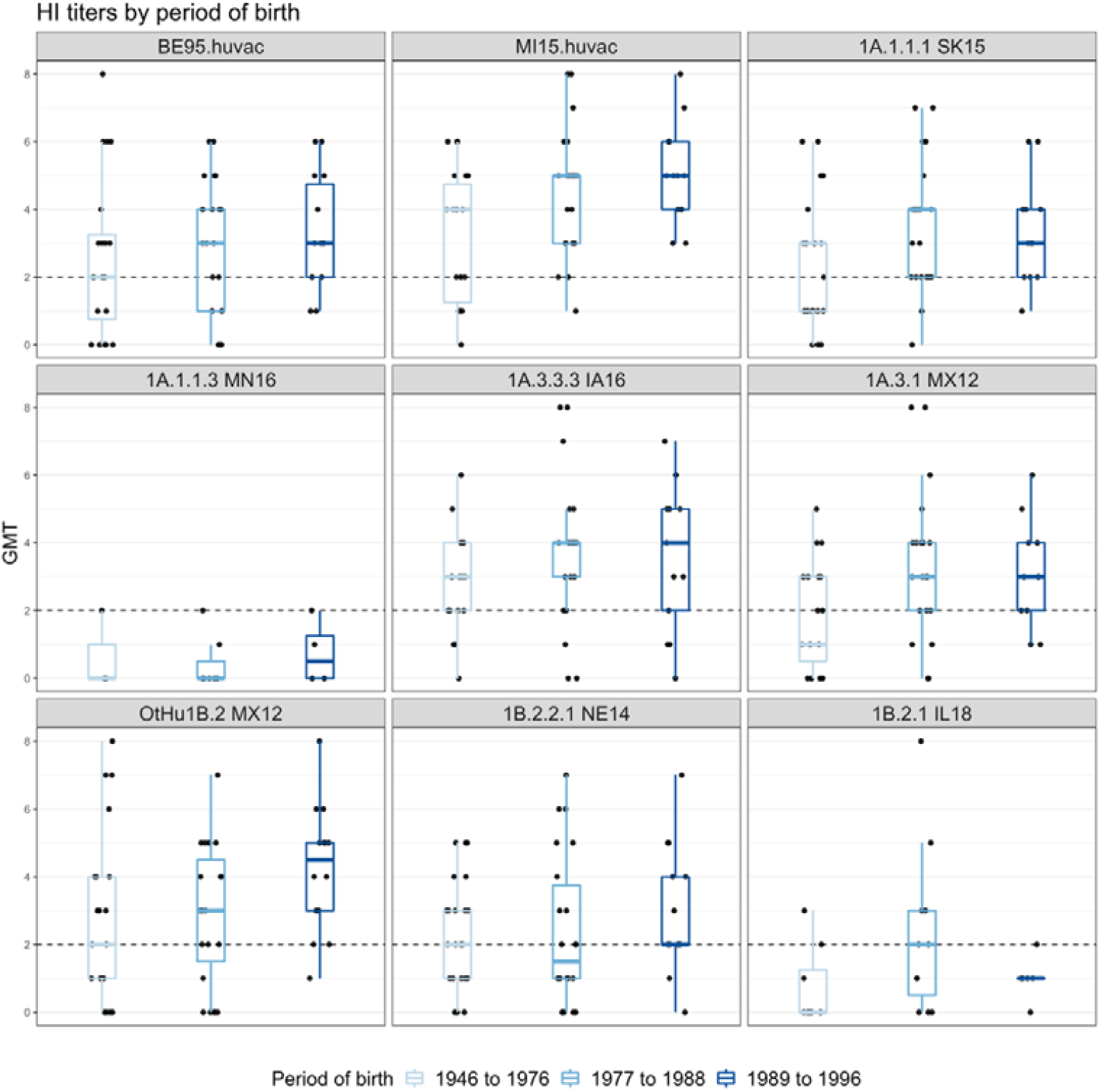
Hemagglutination inhibition responses of post-vaccination and post-exposure human sera stratified by period of birth against North American H1 swine strains. Shades of light to dark blue represent the periods of birth of the human participants (1946 to 1976, 1977 to 1988, and 1989 to 1996). Box plots show the median HI titers against H1 strains with 5th and 95th percentile and standard deviation. Each dot represents the geometric mean titer (GMT) of human sera on the y-axis against each strain (shown on x-axis). The grey dotted line indicates the minimum positive HI titers threshold (≥40 or 2). No major differences are seen between the different age groups.

We hypothesized that the sera that showed high cross-reactivity with the post-pandemic MI15 human vaccine strain, either by exposure or vaccination, would also have higher responses to those swine strains which are antigenically similar to the other 1A lineage strains derived from pre-2009 human seasonal strains (MX12 1A.3.1 (AU 2.99), SK15 1A.1.1.1 (AU 3.05), IA16 1A.3.3.3 (AU 4.29). To address this question, we divided the titers into three groups of high, medium, and low responders to the MI15 and found a modest increase in the titers of high MI15 responders against the swine 1A.x viruses (see **Supplementary Figure S4A)**. This pattern was not found for the distribution of titers for MX12 OtHu1B.2 (AU 8.15) and NE14 1B.2.2.1 (AU 6.38), or the MN16 1A.1.1.3 (AU 8.90) and IL18 1B.2.1 (AU 9.72) strains. There was no effect of age since both high and low responders were found within each age-group (**Supplementary Figure S4B**).

## Discussion

Influenza A viruses in swine have posed a consistent challenge to food and animal security around the world. In this study, we quantified the genetic and antigenic diversity in swine influenza A H1 viruses in Canada, USA, and Mexico isolated between 2012 and 2018. We explicitly defined how evolution in swine following introduction of human seasonal IAV lineages into pigs has resulted in significant drift from both human seasonal vaccines and pandemic preparedness candidate vaccine viruses. Our risk assessment pipeline integrated data derived from genomic surveillance and antigenic characterization incorporated into an objective strain selection process. This process identified priority swine IAV genetic clades for assessment of zoonotic potential using human population sera. This process quantified the public health risk of the genetically and antigenically diverse North American swine H1 influenza viruses and supported a need for rigorous evaluation of IAV at the human-swine interface in risk assessment. Our characterization identified H1 IAVs circulating in swine to which the human population likely has minimal immunity from either prior exposure or current vaccine efforts.

Surveillance for swine IAVs revealed an expansion in the diversity of some genetic clades in North America, notably the 1A.1.1 (alpha) (Nelson *et al*., 2017). A single clade of North American 1A.1.1 viruses was defined in the current global swine H1 classification scheme (Anderson *et al*., 2016). Additional surveillance and our characterization revealed regional persistence and circulation of genetically and antigenically distinct viruses within different statistically supported clades in 1A.1.1. Currently circulating 1A.1.1 strains formed two major groups, both phylogenetically and antigenically. Within this study we defined the group reported as a-1 from Nelson et al. 2017 as 1A.1.1.1, and the group reported as a-3 from Nelson et al. 2017 with a two amino acid deletion in the HA1 defined as 1A.1.1. 3. A third clade, the a-2, was identified in Nelson et al. 2017, with the global name 1A.1.1.2, but they represent a small fraction of detections in Canada, so we did not antigenically characterize these viruses.

The antigenic distances between swine IAV strains and human vaccine strains from the past ∼50 years (1977 – 2015) were heterogeneous. The 1A lineage swine viruses averaged between 5.51 to 8.4 AU from the human vaccine strains. This diversity reflects decades of evolution in swine following the human-to-swine spillover event. The 1B lineage swine viruses, originating from human seasonal strains from the 2000s, remained antigenically similar to the putative human ancestral strains. However, these contemporary swine strains had an average distance of 3.72 to 6.14 AU to more recent H1 human seasonal vaccine strains. These data are consistent with estimates of H1 1B viruses antigenic drift of 0.17-0.85 AU per year (Lewis *et al*., 2016) and reflect introduction and decades of evolution in regional swine populations. Notably, some IAVs, such as the IL18 1B.2.1 virus, were antigenically very distant from all human vaccine strains (average AU 7.31 and 10.37 from post-2000 and post-pandemic human strains respectively).

This antigenic diversity in cross-reactivity characterized using swine and ferret antisera was reflected in the human serology assays, where there was also heterogenous cross-reactivity in the human population samples. Consequently, though some 1A and 1B viruses retain some antigenic cross-reactivity with human vaccine strains, they fall above the threshold that is applied to guide vaccine strain updates and human population immunity will reflect this antigenic variation. In our data, we did not find associations between age, sex, or prior exposure that indicated a demographic of the human population more at risk to an incursion by swine IAV into humans. Instead, our data demonstrated that certain clades of swine IAV represent a more pressing zoonotic threat, and this dynamic is the result of evolution within swine unrelated to human population immunity. Although our cohorts were not comprehensive, they were representative of various ages, genders and two distinct geographies. However, we did not have human sera from a birth year period that would most likely have encountered the progenitors of the swine 1B viruses (early 2000s) in early childhood. Given the spatial heterogeneity of contemporary IAV lineages in pigs might influence the emergent risk at the human-animal interface, future work should focus on regional risk assessments using locally derived human serum cohorts. For example, surveillance efforts in Mexico and Canada suggest distinct evolutionary dynamics in those regions with different continuously circulating lineages in swine herds (Mena *et al*., 2016; Nelson *et al*., 2017). Similarly, assessing zoonotic risk using human sera collected in the US from occupationally exposed humans and from groups at critical swine-human interface settings (i.e., agricultural fairs) may be appropriate. Despite these caveats, our data conclusively reveal that evolution of IAV in swine results in specific clades (1A.1.1.3 and 1B.2.1) across two major lineages to which almost no pre-existing immunity exists in the human population.

A key component in human pandemic preparedness is surveillance for currently circulating and emerging IAVs in human populations around the world. These viruses are genetically and antigenically characterized for human risk assessment using polyclonal ferret antisera raised to human vaccine strains or to CVVs. When detected, cases of variant swine influenza A viruses are characterized against existing CVV and potentially identified for new CVV if not recognized by existing ferret anti-sera (Anderson *et al*., 2020). Our data demonstrate that CVVs address some of the gaps in cross-reactivity from human population sera. However, the antigenic evolution of influenza in swine is dynamic and we have little understanding of the breath of diversity in contemporary global swine influenza viruses as it relates to human immunity. Given the correlation between antigenic distance of swine strains to human vaccine strains and measures of reduced human population immunity, our process presents an efficient way to evaluate swine IAV for zoonotic threat. Linking sustainable in-field genomic surveillance efforts with broader cross-HI assays and more representative cohorts of human sera can establish a global pandemic preparedness protocol.

Our data revealed at least 12 genetic clades from within two major evolutionary H1 lineages cocirculating in North American swine. These genetic clades demonstrated considerable antigenic diversity, and swine strains typically had significant losses in cross reactivity to human vaccines in use from 1977 to 2015. Using a metric that encompassed genetic and antigenic diversity, we objectively ranked H1 swine strains, and tested seven prioritized viruses against two cohorts of human sera to as one indication of zoonotic potential. These data demonstrated significant variation in human antibody recognition of the swine strains among and between phylogenetic clades of swine H1. Two of the swine strains from major H1 clades (1A.1.1.3 and 1B.2.1) did not react with most of the samples of human population sera. The observed diversity of swine IAV represents a significant challenge to pandemic risk assessment, but our process that integrates genomic surveillance, antigenic characterization, and testing against human population sera provides a template for objective risk assessment. Building on the system reported here, subsequent testing against representative human sera from the region where the swine strains circulate and from additional birth years would add valuable information to determining the pandemic potential of IAV in swine. These data are fluid and will need future re-assessment as IAV continues to evolve in swine, but this is the first comprehensive report to identify H1 clades in swine that represent zoonotic threats, informing the design of candidate vaccine viruses for humans and identifying strains that may be preemptively targeted through vaccination in the pig population.

## Acknowledgments

We gratefully acknowledge pork producers, swine veterinarians, and laboratories for participating in the USDA Influenza A Virus in Swine Surveillance System and publicly sharing sequences in NCBI GenBank. We thank Michelle Harland, Gwen Nordholm, and Marcus Bolton for laboratory assistance and Jason Huegel, Keiko Sampson, and Justin Miller for animal care and handling assistance. We thank Susan Detmer from the University of Saskatchewan for swine H1N1 viruses and antisera and Adolfo García-Sastre and Ignacio Mena from the Icahn School of Medicine at Mount Sinai, NY, for providing Mexican swine H1N1 strains. We thank Todd Davis and members of the Virology, Surveillance, and Diagnosis Branch, Influenza Division, Centers for Disease Control and Prevention, for provision of reagents used in these studies. Data and code used in this research are available in a GitHub repository https://github.com/flu-crew/h1-risk-pipeline. This work was supported in part by the U.S. Department of Agriculture (USDA) Agricultural Research Service (ARS project number 5030-32000-231-000D); a National Institute of Allergy and Infectious Diseases (NIAID) at the National Institutes of Health (NIH) Center of Excellence in Influenza Research and Surveillance interagency agreement associated with the Center of Research in Influenza Pathogenesis (HHSN272201400008C) and the Johns Hopkins Center of Excellence in Influenza Research and Surveillance (HHSN272201400007C); the NIAID, NIH, Department of Health and Human Services (Contract No. 75N93021C00015); J.B.K, C.K.S and J.C. were supported by an appointment to the USDA Agricultural Research Service Research Participation Program of the Oak Ridge Institute for Science and Education (ORISE) through an interagency agreement between the U.S. Department of Energy (DOE) and USDA Agricultural Research Service (contract number DE-AC05-06OR23100); and the SCINet project of the USDA Agricultural Research Service (ARS project number 0500-00093-001-00-D). The funders had no role in study design, data collection and interpretation, or the decision to submit the work for publication. Mention of trade names or commercial products in this article is solely for the purpose of providing specific information and does not imply recommendation or endorsement by the USDA, DOE, or ORISE. USDA is an equal opportunity provider and employer.

