## Supplemental files for "Antigenic characterization and pandemic risk assessment of North American H1 influenza A viruses circulating in swine"

**Abstract**

The first pandemic of the 21st century was caused by an H1N1 influenza A virus (IAV) introduced from pigs into humans, highlighting the importance of swine as reservoirs for pandemic viruses. Two major lineages of swine H1 circulate in North America: the 1A classical swine lineage (including the 2009 pandemic H1N1) and 1B human seasonal-like lineage. Here, we investigated the evolution of these H1 IAV lineages in North American swine and their potential pandemic risk. We assessed the antigenic distance between the HA of representative swine H1 and human seasonal vaccine strains (1978-2015) in hemagglutination inhibition (HI) assays using a panel of monovalent anti-sera raised in pigs. Antigenic cross-reactivity varied by strain but was associated with genetic distance. Generally, swine 1A lineage viruses that seeded the 2009 H1 pandemic were antigenically most similar to H1 pandemic vaccine strains, with the exception of viruses in the genetic clade 1A.1.1.3 that had a two-amino acid deletion mutation near the receptor-binding site, dramatically reducing antibody recognition. The swine 1B lineage strains, which arose from previously circulating (pre-2009 pandemic) human seasonal viruses, were more antigenically similar to pre-2009 human seasonal H1 vaccine viruses than post-2009 strains. Human population immunity was measured by cross-reactivity in HI assays to representative swine H1 strains. There was a broad range of titers against each swine strain that was not associated with age, sex, or location. However, there was almost no cross-reactivity in human sera to the 1A.1.1.3 and 1B.2.1 genetic clades of swine viruses, and the 1A.1.1.3 and 1B.2.1 clades were also the most antigenically distant from all human vaccine strains. Our data demonstrate that antigenic distances of representative swine strains from human vaccine strains represent a rational assessment of swine IAV for zoonotic risk research and pandemic preparedness prioritization.

### Supplementary information

S1

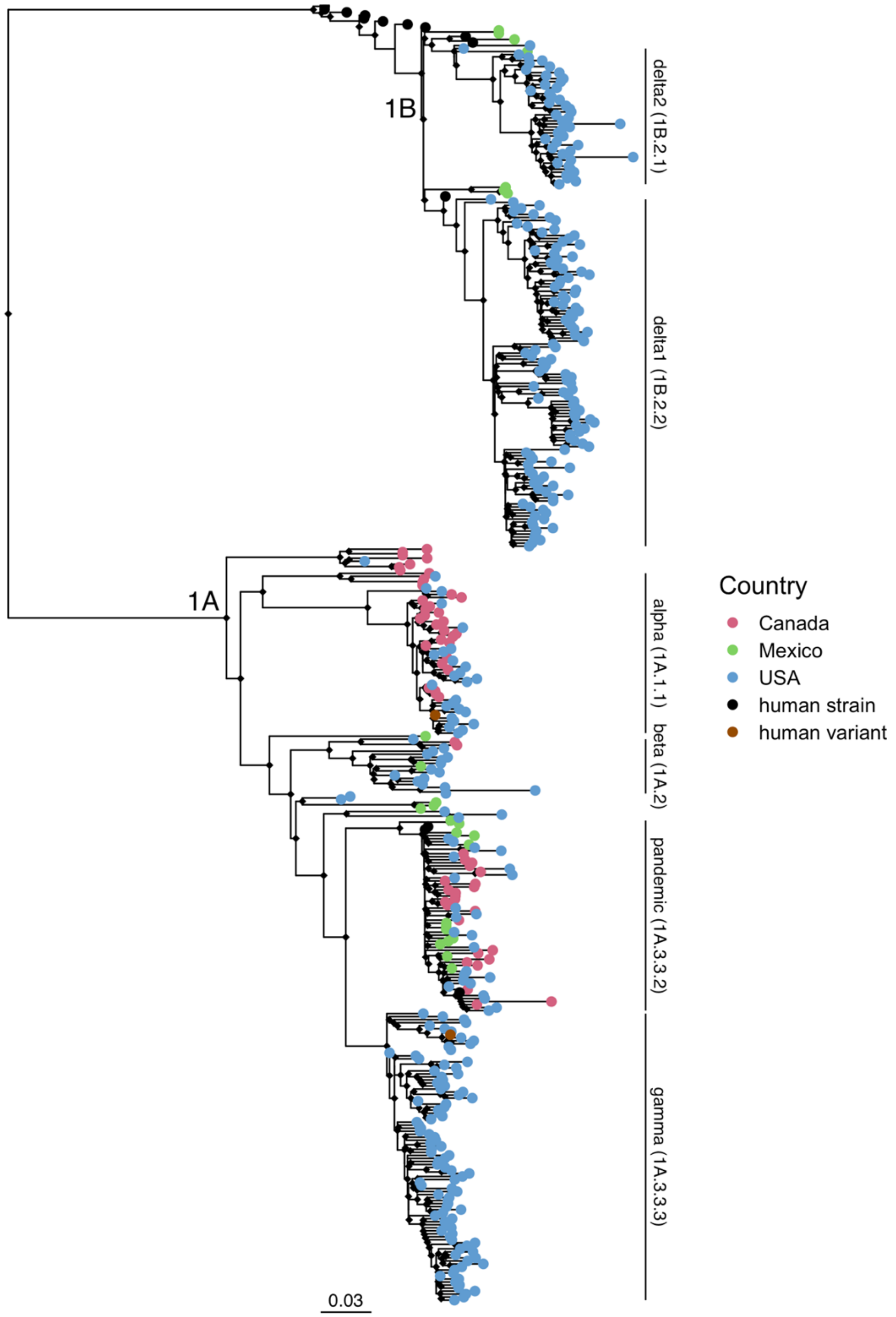


**Figure S1.** Maximum-likelihood phylogenetic tree of swine influenza A viruses isolated in North America between 2012-2019 and human vaccine strains. Tips are coloured by country of origin of swine strains. Human strains are in black and swine-to-human transmission cases are in brown.

S2

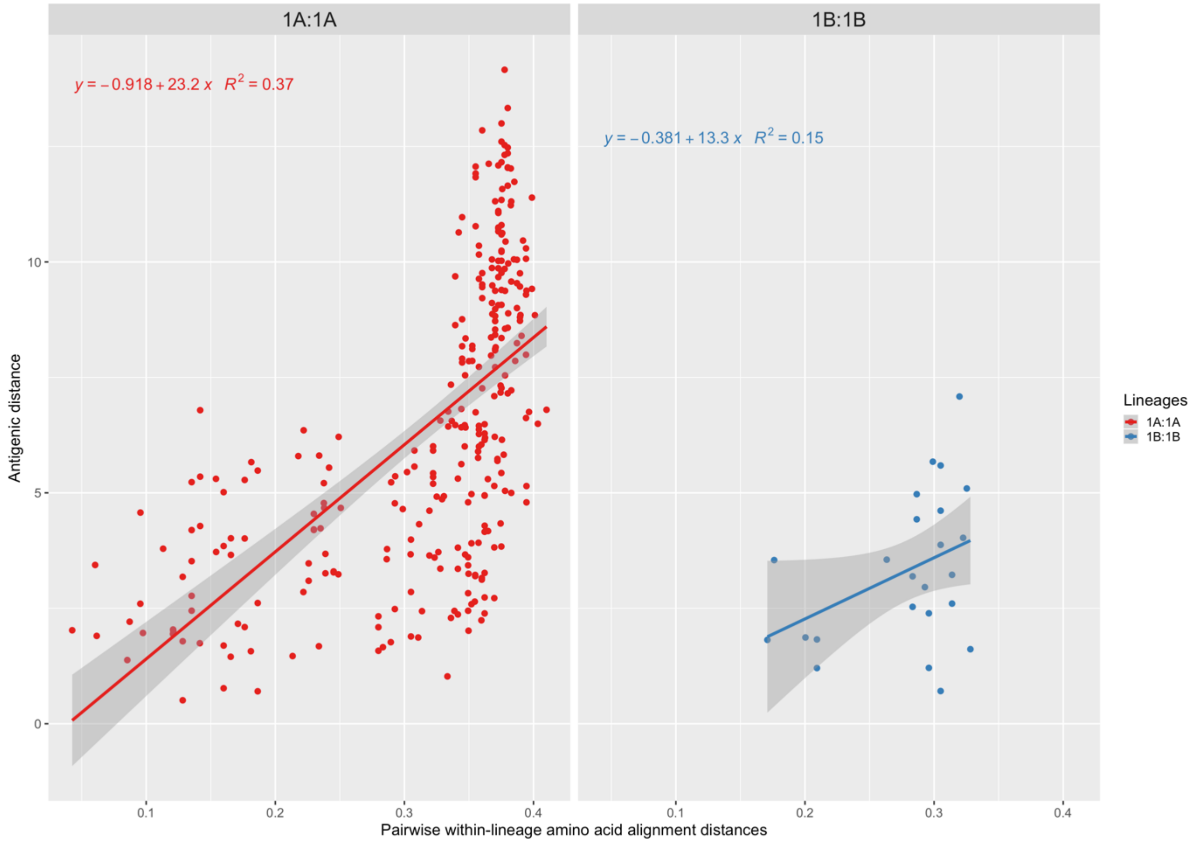


**Figure S2.** A scatterplot of genetic (amino acid) versus antigenic distance between viruses within the 1A and 1B lineages with a linear model regression line fitted to the data.

S3

A

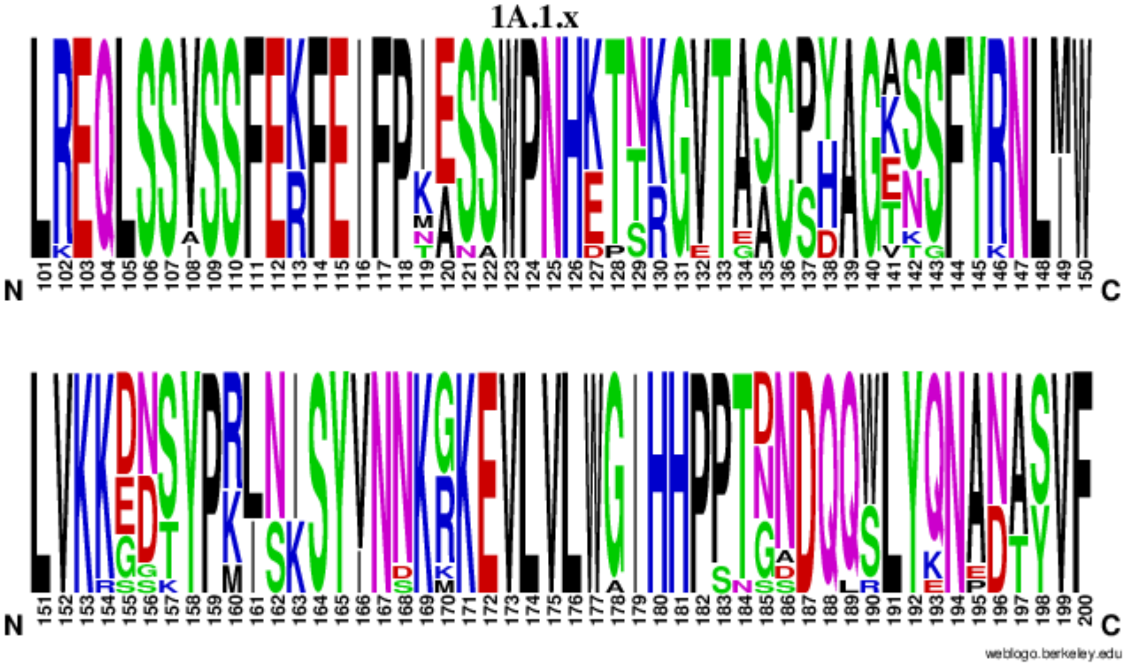


B

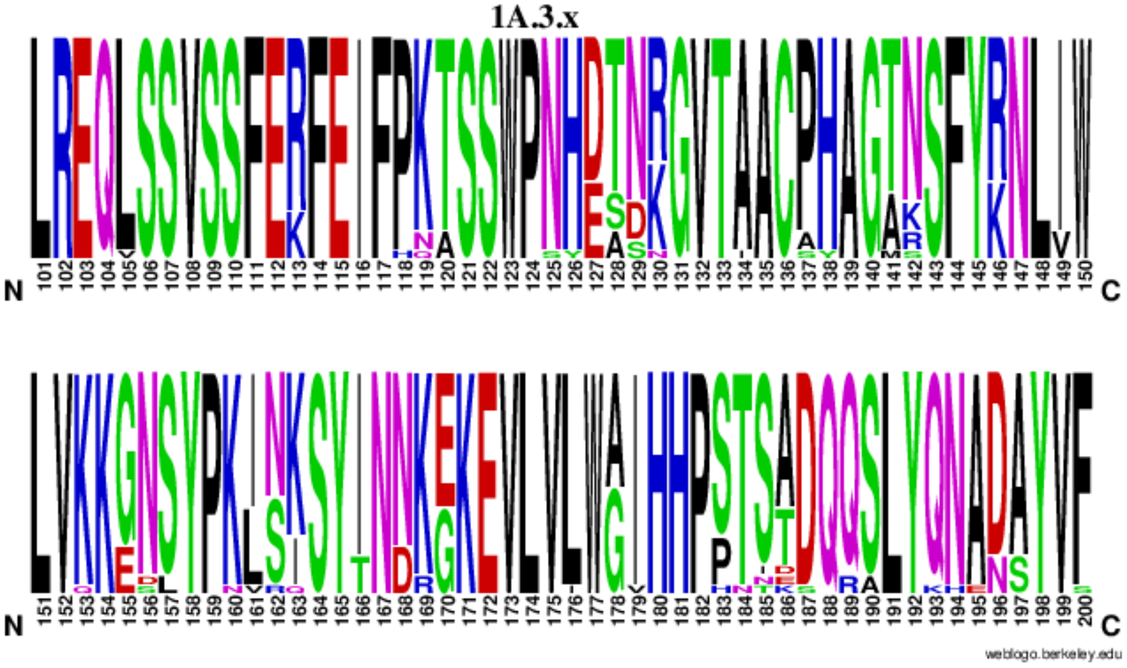

C

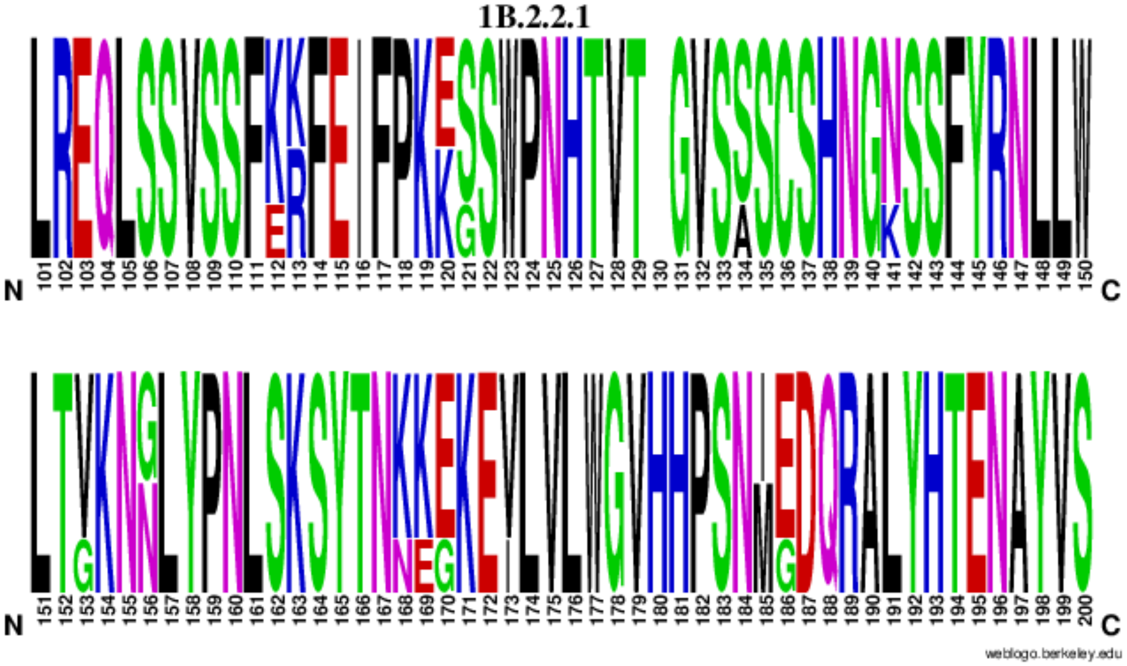


D

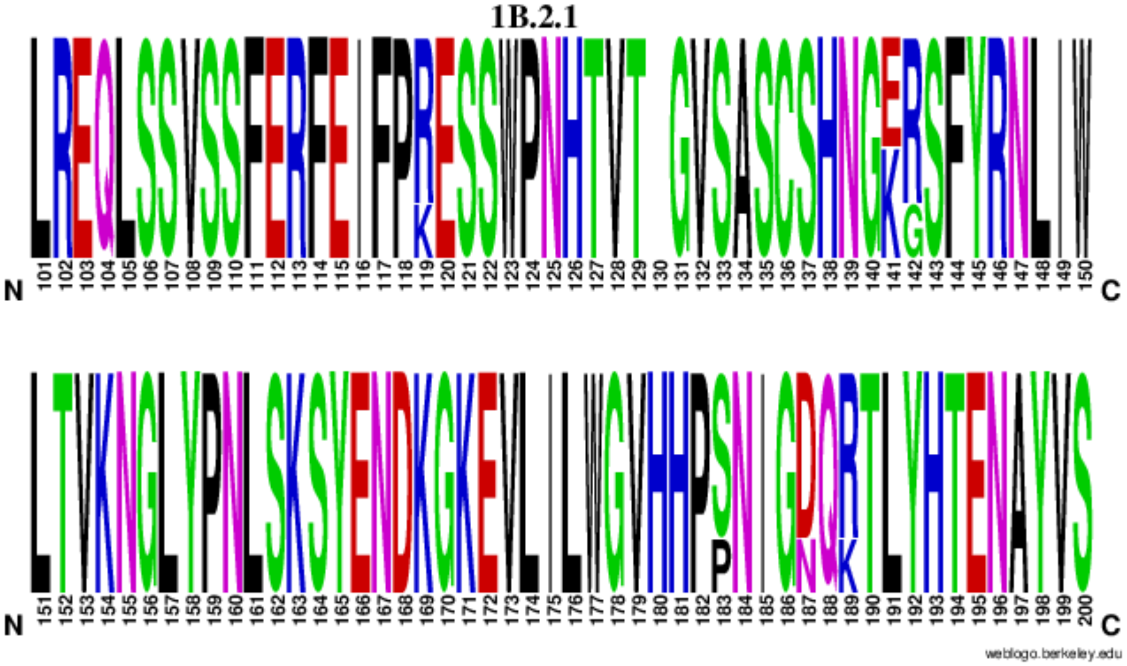


**Figure S3 (A-D).** Web-logo of the amino acid alignment (101-200) of each sub-lineage of North American swine influenza A virus: A (1A.1.x), B (1A.3.x), C (1B.2.1) and D (1B.2.2.1). Logo shows variation at each amino acid residue in a region of the protein in which largely determines antigenicity (recognised by host antibodies). (Crooks et al. 2004)

S4

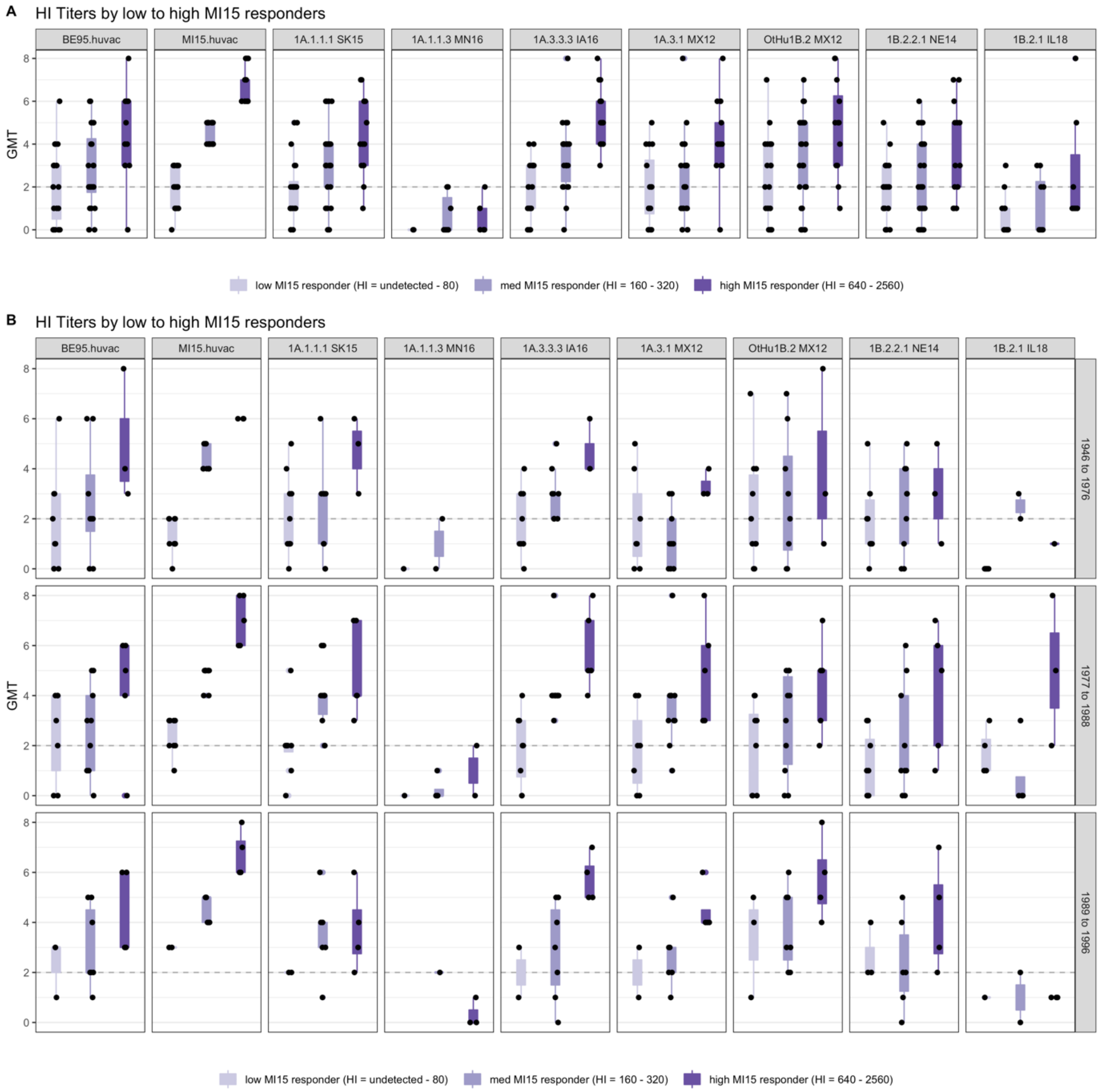


**Figure S4 (A-B).** Combined post-exposure and post-vaccination titres of human sera against representative swine strains, stratified by (A) low – to – high responders to MI15 (A/Michigan/45/2015) vaccine strain, (B) low – to – high responders to MI15 and period of birth (1945-76, 1977-88, 1989-96). Violin box plots show the median of aggregated HI titres against H1N1 strains with 5th and 95th percentile and standard deviation. Each dot represents the GMT: geometric mean titre, log2 (HI titre /10) of human sera on the y-axis against each strain (shown on x-axis). Cohorts are combined as results are similar for individual cohorts).

S5

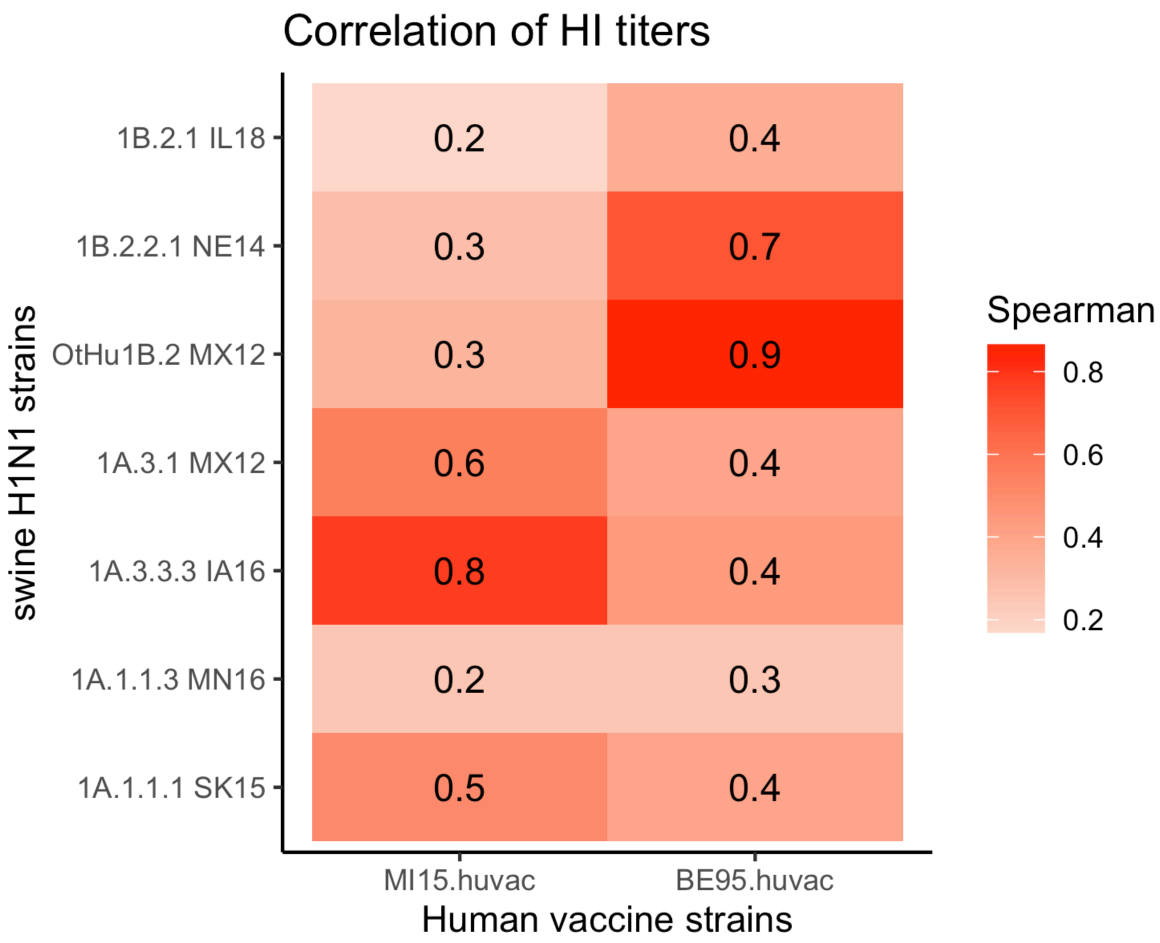


**Figure S5.** Spearman's correlation between HI titers against North American H1 swine strains and human seasonal H1 vaccine strains of post-exposure convalescent and post-vaccination human sera cohorts (combined as results similar to individual cohorts).

**Table S1.** Swine and human influenza A viruses and sera raised against them used in this study.

| Viruses tested agianst swine sera | virus clade | virus subtype | HA reference |
|---|---|---|---|
| A/swine/Alberta/SD0125/2015 | 1A.1.1 | H1N1 | MF768555 |
| A/swine/Saskatchewan/SD0094/2015* | 1A.1.1 | H1N1 | MF768483 |
| A/swine/Alberta/SD0014/2013 | 1A.1.1 | H1N1 | CY195078 |
| A/swine/Alberta/SD0154/2016 | 1A.1.1 | H1N1 | MF768523 |
| A/swine/Manitoba/D0333/2014 | 1A.1.1 | H1N2 | CY195575 |
| A/swine/Manitoba/D0392/2015 | 1A.1.1 | H1N2 | MF768531 |
| A/swine/Saskatchewan/SD0102/2015 | 1A.1.1 | H1N2 | MF768539 |
| A/swine/Manitoba/SD0114/2015 | 1A.1.1 | H1N2 | MF768507 |
| A/swine/Alberta/SD0191/2016 | 1A.1.1 | H1N2 | MF768475 |
| A/swine/Saskatchewan/SD0200/2016 | 1A.1.1 | H1N2 | MF768547 |
| A/swine/Manitoba/D0348/2014 | 1A.1.1 | H1N2 | CY195295 |
| A/Minnesota/45/2016 | 1A.1.1 | H1N2 | EPI760602 |
| A/swine/Minnesota/A01781045/2016* | 1A.1.1 | H1N2 | KX928680 |
| A/swine/Mexico/AVX23/2012* | 1A.2-3-like | H1N1 | KU976649 |
| A/swine/Illinois/A01493472/2014 | 1A.3.3.2 | H1N1 | KJ701784 |
| A/California/04/2009 | 1A.3.3.2 | H1N1 | GQ280797 |
| A/Mexico/4108/2009 | 1A.3.3.2 | H1N1 | GQ223112 |
| A/Michigan/45/2015* | 1A.3.3.2 | H1N1 | KU933493 |
| A/swine/Mexico/AVX31/2012 | 1A.3.3.2 | H1N1 | KU976950 |
| A/swine/Mexico/AVX44/2012 | 1A.3.3.2 | H1N1 | KU976796 |
| A/swine/North Carolina/A02076926/2015 | 1A.3.3.3 | H1N1 | KT429540 |
| A/swine/North Carolina/A01841602/2015 | 1A.3.3.3 | H1N1 | KR088267 |
| A/swine/Ohio/A01847657/2015 | 1A.3.3.3 | H1N1 | KR780630 |
| A/Ohio/09/2015 | 1A.3.3.3 | H1N1 | EPI587643 |
| A/swine/Minnesota/A01567490/2014 | 1A.3.3.3 | H1N1 | KP662638 |
| A/swine/North Carolina/A01797415/2015 | 1A.3.3.3 | H1N1 | KU357008 |
| A/swine/Iowa/A01731653/2016* | 1A.3.3.3 | H1N1 | KU877398 |
| A/swine/North Carolina/A01730369/2016 | 1A.3.3.3 | H1N1 | KU695684 |
| A/swine/Oklahoma/A01409770/2014 | 1B.2.1 | H1N2 | KJ437589 |
| A/swine/Illinois/A02139356/2018* | 1B.2.1 | H1N2 | MG917068 |
| A/swine/Michigan/A01104117/2018 | 1B.2.1 | H1N2 | MH758776 |
| A/swine/Minnesota/A01134353/2011 | 1B.2.2.1 | H1N2 | JQ906881 |
| A/swine/Nebraska/A01492366/2014* | 1B.2.2.1 | H1N2 | KJ549771 |
| A/swine/Illinois/A01644323/2018 | 1B.2.2.1 | H1N2 | MG825101 |
| A/Brazil/11/1978 | Other-Human | H1N1 | CY020293 |
| A/New Caledonia/20/1999 | Other-Human-1B.2 | H1N1 | EU103824 |
| A/swine/Mexico/AVX18/2012* | Other-Human-1B.2 | H1N2 | KU976609 |
| A/swine/Mexico/AVX61/2013 | Other-Human-1B.2 | H1N2 | KU976596 |
| A/Solomon Islands/3/2006 | Other-Human-1B.2 | H1N1 | EU124135 |
| A/Singapore/6/1986 | Other-Human-1B.2 | H1N1 | CY020477 |
| A/Taiwan/1/1986 | Other-Human-1B.2 | H1N1 | X17224 |
| A/Texas/36/1991 | Other-Human-1B.2 | H1N1 | AJ457908 |
| A/Beijing/262/1995* | Other-Human-1B.2 | H1N1 | AJ457900 |
| A/Michigan/2/2003 | Other-Human-1B.2 | H1N2 | CY016324 |
| A/Brisbane/59/2007 | Other-Human-1B.2 | H1N1 | KF009550 |

*viruses tested against human sera

| Swine sera raised against virus | virus clade | virus subtype | HA reference |
|---|---|---|---|
| A/swine/Iowa/1973 | 1A.1.1 | H1N1 | EU139826 |
| A/swine/Alberta/SD0125/2015 | 1A.1.1 | H1N1 | MF768555 |
| A/swine/Alberta/SD0014/2013 | 1A.1.1 | H1N1 | CY195078 |
| A/swine/Alberta/SD0154/2016 | 1A.1.1 | H1N1 | MF768523 |
| A/swine/Manitoba/D0333/2014 | 1A.1.1 | H1N2 | CY195575 |
| A/swine/Minnesota/02053/2008 | 1A.1.1 | H1N1 | HM461762 |
| A/swine/Manitoba/D0392/2015 | 1A.1.1 | H1N2 | MF768531 |
| A/swine/Saskatchewan/SD0102/2015 | 1A.1.1 | H1N2 | MF768539 |
| A/swine/Manitoba/SD0114/2015 | 1A.1.1 | H1N2 | MF768507 |
| A/swine/Saskatchewan/SD0142/2016 | 1A.1.1 | H1N2 | MK462362 |
| A/swine/Minnesota/A01781045/2016 | 1A.1.1 | H1N2 | KX928680 |
| A/swine/Minnesota/37866/1999 | 1A.2-3-like | H1N1 | EU139827 |
| A/swine/South Dakota/A01349306/2013 | 1A.3.2 | H1N1 | KC844200 |
| A/swine/Minnesota/1192/2001 | 1A.3.3-like | H1N2 | EU139828 |
| A/Mexico/4108/2009 | 1A.3.3.2 | H1N1 | GQ223112 |
| A/swine/Illinois/A01493472/2014 | 1A.3.3.2 | H1N1 | KJ701784 |
| A/California/04/2009 | 1A.3.3.2 | H1N1 | GQ280797 |
| A/Michigan/45/2015 | 1A.3.3.2 | H1N1 | KU933493 |
| A/swine/Minnesota/00194/2003 | 1A.3.3.3 | H1N2 | EU139830 |
| A/swine/Kansas/00246/2004 | 1A.3.3.3 | H1N2 | CY081680 |
| A/swine/OH/511445/2007 | 1A.3.3.3 | H1N1 | EU604689 |
| A/swine/Missouri/02060/2008 | 1A.3.3.3 | H1N1 | CY082655 |
| A/swine/North Carolina/02023/2008 | 1A.3.3.3 | H1N1 | HM461818 |
| A/swine/Ohio/02026/2008 | 1A.3.3.3 | H1N1 | CY082631 |
| A/Ohio/09/2015 | 1A.3.3.3 | H1N1 | EPI587643 |
| A/swine/Iowa/A01731653/2016 | 1A.3.3.3 | H1N1 | KU877398 |
| A/swine/Illinois/00685/2005 | 1B.2.1 | H1 | CY081899 |
| A/swine/Oklahoma/A01409770/2014 | 1B.2.1 | H1N2 | KJ437589 |
| A/swine/Minnesota/A01134353/2011 | 1B.2.2.1 | H1N2 | JQ906881 |
| A/swine/Nebraska/A01492366/2014 | 1B.2.2.1 | H1N2 | KJ549771 |
| A/Brazil/11/1978 | Other-Human | H1N1 | CY020293 |
| A/Taiwan/1/1986 | Other-Human-1B.2 | H1N1 | X17224 |
| A/Beijing/262/1995 | Other-Human-1B.2 | H1N1 | AJ457900 |
| A/New Caledonia/20/1999 | Other-Human-1B.2 | H1N1 | EU103824 |
| A/Solomon Islands/3/2006 | Other-Human-1B.2 | H1N1 | EU124135 |

**Table S2.** Hemagglutination inhibition titers against homologous and heterologous strains extracted from the antigenic map.

| Antigen names | Swine serum IDs | EXP428-P592 | EXP428-P560 | EXP428-P569 | EXP428-P578 | EXP428-P581 | EXP467-P821 | EXP569-P859 | EXP569-P860 | EXP480-P868 | EXP480-P869 | EXP480-P864 | EXP480-P865 | EXP480-P866 | EXP480-P867 | EXP569B-P927 | EXP569B-P928 | EXP496M-P960 | EXP496M-P962 | EXP496M-963 | EXP496M-966 | EXP604-P50 | EXP604-P51 | EXP592B-P3 |
|---|---|---|---|---|---|---|---|---|---|---|---|---|---|---|---|---|---|---|---|---|---|---|---|---|
| | Passage/Serum names | A/SWINE/MINNESOTA/1192/2001 | A/SWINE/MINNESOTA/00194/2003 | A/SWINE/MINNESOTA/00194/2003 | A/SWINE/KANSAS/00246/2004 | A/SWINE/KANSAS/00246/2004 | A/SWINE/OHIO/511445/2007 | A/SWINE/OHIO/511445/2007 | A/SWINE/OHIO/511445/2007 | A/SWINE/MISSOURI/02060/2008 | A/SWINE/MISSOURI/02060/2008 | A/SWINE/NORTH_CAROLINA/02023/2008 | A/SWINE/NORTH_CAROLINA/02023/2008 | A/SWINE/OHIO/02026/2008 | A/SWINE/OHIO/02026/2008 | A/SWINE/SOUTH_DAKOTA/A01349306/2013 | A/SWINE/SOUTH_DAKOTA/A01349306/2013 | A/CALIFORNIA/4/2009 | A/CALIFORNIA/4/2009 | A/MEXICO/4108/2009 | A/MEXICO/4108/2009 | A/OHIO/9/2015 | A/OHIO/9/2015 | A/SWINE/ILLINOIS/A01493472/2014 |
| A/SWINE/NORTH_CAROLINA/A02076926/2015 | MDCK2 | 80 | <10 | 40 | 80 | 80 | 80 | 80 | 40 | 80 | 80 | <10 | 40 | <10 | 320 | 40 | 20 | 640 | 640 | 160 | 80 | 20 | 80 | 20 |
| A/SWINE/NORTH_CAROLINA/A01841602/2015 | MDCK2 | <10 | <10 | <10 | 10 | 40 | 40 | 10 | 40 | 80 | 40 | <10 | 20 | <10 | 160 | 20 | <10 | 80 | 80 | 80 | 20 | 20 | 40 | 10 |
| A/SWINE/OHIO/A01847657/2015 | MDCK2 | <10 | <10 | <10 | <10 | 40 | 20 | <10 | 20 | 20 | 20 | <10 | 10 | <10 | 160 | 40 | <10 | 80 | 80 | 40 | 20 | 20 | 40 | <10 |
| A/OHIO/9/2015 | MDCK2 | 20 | <10 | <10 | 20 | 40 | 40 | <40 | * | 160 | 80 | <10 | 10 | <10 | 160 | 57 | <20 | 127 | 160 | 80 | 80 | 127 | 160 | 10 |
| A/SWINE/ILLINOIS/A01493472/2014 | MDCK2 | 320 | 320 | 320 | 320 | 160 | 80 | 160 | 80 | 80 | 160 | 40 | 80 | 10 | 40 | 320 | 80 | 2560 | 5120 | 2560 | 2560 | <10 | <10 | 160 |
| A/SWINE/MINNESOTA/A01567490/2014 | MDCK2 | 160 | 320 | 160 | 1280 | 320 | 1280 | 640 | 160 | 640 | 640 | 160 | 320 | 80 | 640 | 160 | 80 | 1280 | 1280 | 320 | 320 | 20 | 40 | <10 |
| A/CALIFORNIA/4/2009 | MDCK4 | 160 | 160 | 160 | 80 | 160 | 80 | 101 | * | 20 | 80 | 40 | 40 | 10 | 40 | 226 | 113 | 3225 | 5120 | 2560 | 2560 | <10 | <10 | 80 |
| A/MEXICO/4108/2009 | MDCK3 | 160 | 320 | 160 | 160 | 160 | 160 | 160 | 80 | 160 | 320 | 40 | 80 | 20 | 80 | 640 | 80 | 5120 | 5120 | 2560 | 1280 | 10 | <10 | 80 |
| A/BRAZIL/11/1978 | MDCK2 | * | * | * | * | * | * | <10 | <10 | * | * | * | * | * | * | <10 | <10 | <10 | <10 | * | * | <10 | <10 | * |
| A/NEW_CALEDONIA/20/1999 | E2 | * | * | * | * | * | * | <10 | <10 | * | * | * | * | * | * | 80 | 20 | <20 | <40 | * | * | <20 | <10 | * |
| A/MICHIGAN/45/2015 | MDCK3 | * | * | * | * | * | * | * | * | * | * | * | * | * | * | 80 | 40 | * | * | * | * | <10 | <10 | * |
| A/SWINE/NORTH_CAROLINA/A01797415/2015 | MDCK3 | * | * | * | * | * | * | 640 | 640 | * | * | * | * | * | * | 160 | 40 | 640 | 640 | * | * | 40 | 40 | * |
| A/SWINE/IOWA/A01731653/2016 | MDCK3 | * | * | * | * | * | * | 80 | 80 | * | * | * | * | * | * | 80 | 20 | 160 | 320 | * | * | <10 | 10 | * |
| A/SWINE/MEXICO/AVX31/2012 | MDCK2 | * | * | * | * | * | * | 160 | 160 | * | * | * | * | * | * | 640 | 320 | 2560 | 2560 | * | * | 20 | 10 | * |
| A/SWINE/MEXICO/AVX23/2012 | MDCK2 | * | * | * | * | * | * | 160 | 160 | * | * | * | * | * | * | 40 | 10 | 320 | 640 | * | * | <10 | <10 | * |
| A/SWINE/MEXICO/AVX44/2012 | MDCK2 | * | * | * | * | * | * | <10 | <10 | * | * | * | * | * | * | 640 | 80 | 80 | 160 | * | * | <10 | <10 | * |
| A/SWINE/MEXICO/AVX18/2012 | MDCK2 | * | * | * | * | * | * | <10 | <10 | * | * | * | * | * | * | <10 | <10 | <10 | <10 | * | * | 20 | <10 | * |
| A/SWINE/MEXICO/AVX61/2013 | MDCK2 | * | * | * | * | * | * | <10 | <10 | * | * | * | * | * | * | 10 | 10 | <10 | <10 | * | * | 10 | 10 | * |
| A/SOLOMON_ISLANDS/3/2006 | E2 | * | * | * | * | * | * | <10 | <10 | * | * | * | * | * | * | * | * | <10 | <10 | * | * | <10 | <10 | * |
| A/SWINE/ALBERTA/SD0125/2015 | MDCK2 | * | * | * | * | * | * | * | * | * | * | * | * | * | * | * | * | 160 | 320 | * | * | * | * | * |
| A/SWINE/SASKATCHEWAN/SD0094/2015 | MDCK2 | * | * | * | * | * | * | * | * | * | * | * | * | * | * | * | * | 160 | 160 | * | * | * | * | * |
| A/SWINE/ALBERTA/SD0014/2013 | MDCK2 | * | * | * | * | * | * | * | * | * | * | * | * | * | * | * | * | 80 | 160 | * | * | * | * | * |
| A/SWINE/ALBERTA/SD0154/2016 | MDCK2 | * | * | * | * | * | * | * | * | * | * | * | * | * | * | * | * | 320 | 320 | * | * | * | * | * |
| A/SWINE/MANITOBA/D0333/2014 | MDCK2 | * | * | * | * | * | * | * | * | * | * | * | * | * | * | * | * | 40 | 80 | * | * | * | * | * |
| A/SWINE/MANITOBA/D0392/2014 | MDCK2 | * | * | * | * | * | * | * | * | * | * | * | * | * | * | * | * | <10 | <10 | * | * | * | * | * |
| A/SWINE/SASKATCHEWAN/SD0102/2015 | MDCK2 | * | * | * | * | * | * | * | * | * | * | * | * | * | * | * | * | <10 | <10 | * | * | * | * | * |
| A/SWINE/MANITOBA/SD0114/2015 | MDCK2 | * | * | * | * | * | * | * | * | * | * | * | * | * | * | * | * | <10 | <10 | * | * | * | * | * |
| A/SWINE/ALBERTA/SD0191/2016 | MDCK2 | * | * | * | * | * | * | * | * | * | * | * | * | * | * | * | * | <10 | <10 | * | * | * | * | * |
| A/SWINE/SASKATCHEWAN/SD0200/2016 | MDCK2 | * | * | * | * | * | * | * | * | * | * | * | * | * | * | * | * | <10 | <10 | * | * | * | * | * |
| A/SWINE/MANITOBA/D0348/2014 | MDCK2 | * | * | * | * | * | * | * | * | * | * | * | * | * | * | * | * | <10 | <10 | * | * | * | * | * |
| A/MINNESOTA/45/2016 | MDCK3 | * | * | * | * | * | * | * | * | * | * | * | * | * | * | * | * | <10 | <10 | * | * | * | * | * |
| A/SWINE/MINNESOTA/A01781045/2016 | MDCK2 | * | * | * | * | * | * | * | * | * | * | * | * | * | * | * | * | * | * | * | * | * | * | * |
| A/SINGAPORE/1986 | MDCK2 | * | * | * | * | * | * | <10 | <10 | * | * | * | * | * | * | * | * | <10 | 10 | * | * | 10 | 10 | * |
| A/TAIWAN/1/1986 | MDCK2 | * | * | * | * | * | * | <10 | <10 | * | * | * | * | * | * | * | * | <10 | 20 | * | * | <10 | <10 | * |
| A/TEXAS/36/1991 | MDCK2 | * | * | * | * | * | * | <10 | <10 | * | * | * | * | * | * | * | * | 10 | 40 | * | * | <10 | <10 | * |
| A/BEIJING/262/1995 | MDCK2 | * | * | * | * | * | * | <10 | <10 | * | * | * | * | * | * | * | * | <10 | <10 | * | * | <10 | <10 | * |
| A/MICHIGAN/2/2003 | MDCK2 | * | * | * | * | * | * | <10 | <10 | * | * | * | * | * | * | * | * | <10 | <10 | * | * | <10 | <10 | * |
| A/BRISBANE/59/2007 | MDCK3 | * | * | * | * | * | * | 10 | 20 | * | * | * | * | * | * | * | * | <10 | <10 | * | * | 10 | 10 | * |
| A/SWINE/NORTH_CAROLINA/A01730369/2016 | MDCK3 | * | * | * | * | * | * | <10 | <10 | * | * | * | * | * | * | * | * | 160 | 320 | * | * | <10 | <10 | * |
| A/SOLOMON_ISLANDS/3/2006 | E2/M1 | * | * | * | * | * | * | * | * | * | * | * | * | * | * | * | * | * | * | * | * | * | * | * |
| A/SWINE/MINNESOTA/A01134353/2011 | MDCK3 | * | * | * | * | * | * | * | * | * | * | * | * | * | * | * | * | * | * | * | * | * | * | * |
| A/SWINE/NEBRASKA/A01492366/2014 | MDCK3 | * | * | * | * | * | * | * | * | * | * | * | * | * | * | * | * | * | * | * | * | * | * | * |
| A/SWINE/ILLINOIS/A01644323/2018 | MDCK3 | * | * | * | * | * | * | * | * | * | * | * | * | * | * | * | * | * | * | * | * | * | * | * |
| A/SWINE/OKLAHOMA/A01409770/2014 | MDCK3 | * | * | * | * | * | * | * | * | * | * | * | * | * | * | * | * | * | * | * | * | * | * | * |
| A/SWINE/ILLINOIS/A02139356/2018 | MDCK3 | * | * | * | * | * | * | * | * | * | * | * | * | * | * | * | * | * | * | * | * | * | * | * |
| A/SWINE/MICHIGAN/A01104117/2018 | MDCK2 | * | * | * | * | * | * | * | * | * | * | * | * | * | * | * | * | * | * | * | * | * | * | * |

| Antigen names | Swine serum IDs | EXP604-P18 | EXP604-P19 | EXP604-P38 | EXP604-P39 | EXP604-P16 | EXP604-P17 | EXP527-P460 | EXP527-P462 | EXP527-P468 | EXP527-P471 | FLU17-P706 | FLU17-P707 | FLU14-P703 | FLU14-P705 | EXP428-P593 | EXP428-P595 | EXP428-P532 | EXP428-P588 | FLU7-P501 | FLU7-P502 | EXP535-P171 | EXP535-P172 |
|---|---|---|---|---|---|---|---|---|---|---|---|---|---|---|---|---|---|---|---|---|---|---|---|
| | Passage/Serum names | A/BRAZIL/11/1978 | A/BRAZIL/11/1978 | A/TAIWAN/1/1986 | A/TAIWAN/1/1986 | A/BEIJING/262/1995 | A/BEIJING/262/1995 | A/NEW_CALEDONIA/20/1999 | A/NEW_CALEDONIA/20/1999 | A/SOLOMON_ISLANDS/3/2006 | A/SOLOMON_ISLANDS/3/2006 | A/CALIFORNIA/4/2009 | A/CALIFORNIA/4/2009 | A/MICHIGAN/45/2015 | A/MICHIGAN/45/2015 | A/SWINE/IOWA/1973 | A/SWINE/IOWA/1973 | A/SWINE/MINNESOTA/37866/1999 | A/SWINE/MINNESOTA/37866/1999 | A/SWINE/IOWA/A01731653/2016 | A/SWINE/IOWA/A01731653/2016 | A/SWINE/ILLINOIS/00685/2005 | A/SWINE/ILLINOIS/00685/2005 |
| A/SWINE/NORTH_CAROLINA/A02076926/2015 | MDCK2 | * | * | * | * | * | * | * | * | * | * | * | * | * | * | * | * | * | * | * | * | * | * |
| A/SWINE/NORTH_CAROLINA/A01841602/2015 | MDCK2 | * | * | * | * | * | * | * | * | * | * | * | * | * | * | * | * | * | * | * | * | * | * |
| A/SWINE/OHIO/A01847657/2015 | MDCK2 | * | * | * | * | * | * | * | * | * | * | * | * | * | * | * | * | * | * | * | * | * | * |
| A/OHIO/9/2015 | MDCK2 | <10 | <10 | <10 | <10 | <10 | <10 | <10 | <10 | <10 | <10 | 160 | 160 | <10 | 10 | 80 | 160 | 80 | 226 | <10 | <10 | <10 | <10 |
| A/SWINE/ILLINOIS/A01493472/2014 | MDCK2 | * | * | * | * | * | * | * | * | * | * | * | * | * | * | * | * | * | * | * | * | * | * |
| A/SWINE/MINNESOTA/A01567490/2014 | MDCK2 | * | * | * | * | * | * | * | * | * | * | * | * | * | * | * | * | * | * | * | * | * | * |
| A/CALIFORNIA/4/2009 | MDCK4 | <10 | <10 | <10 | <10 | <10 | <10 | <10 | <10 | <10 | <10 | 2560 | 640 | 160 | 160 | 20 | 80 | 80 | 320 | <10 | <10 | <10 | <10 |
| A/MEXICO/4108/2009 | MDCK3 | * | * | * | * | * | * | * | * | * | * | * | * | * | * | * | * | * | * | * | * | * | * |
| A/BRAZIL/11/1978 | MDCK2 | 202 | 320 | 24 | 17 | <40 | <40 | <10 | <10 | <10 | <10 | * | * | <10 | <10 | <10 | <10 | <10 | <10 | <10 | <10 | <10 | <10 |
| A/NEW_CALEDONIA/20/1999 | E2 | <10 | <10 | <20 | <10 | 20 | 28 | 127 | 127 | 25 | 101 | <10 | <10 | <10 | <10 | <10 | <10 | <10 | <10 | <10 | <10 | 101 | 160 |
| A/MICHIGAN/45/2015 | MDCK3 | <10 | <10 | <10 | <10 | <10 | <10 | <10 | <10 | <10 | <10 | 320 | 80 | 160 | 160 | * | * | 40 | 160 | <10 | <10 | * | * |
| A/SWINE/NORTH_CAROLINA/A01797415/2015 | MDCK3 | <10 | <10 | 10 | 20 | 10 | <10 | <10 | <10 | 10 | 40 | 80 | 40 | 20 | 10 | 40 | 160 | 320 | 640 | 320 | 640 | 10 | 20 |
| A/SWINE/IOWA/A01731653/2016 | MDCK3 | <10 | <10 | <10 | 10 | <10 | <10 | <10 | <10 | <10 | <10 | 80 | 20 | 40 | 40 | 20 | 80 | 160 | 320 | 1280 | 640 | <10 | <10 |
| A/SWINE/MEXICO/AVX31/2012 | MDCK2 | 10 | 10 | <10 | <10 | <10 | <10 | <10 | <10 | <10 | <10 | 1280 | 320 | 160 | 160 | 20 | 40 | 320 | 320 | 40 | 20 | <10 | <10 |
| A/SWINE/MEXICO/AVX23/2012 | MDCK2 | <10 | <10 | <10 | <10 | <10 | <10 | <10 | <10 | <10 | <10 | 160 | 160 | 160 | 40 | 20 | 10 | 320 | 640 | 80 | 40 | <10 | <10 |
| A/SWINE/MEXICO/AVX44/2012 | MDCK2 | <10 | <10 | <10 | <10 | <10 | <10 | <10 | <10 | <10 | <10 | 160 | 80 | 40 | 20 | 40 | 40 | 10 | 40 | <10 | <10 | <10 | <10 |
| A/SWINE/MEXICO/AVX18/2012 | MDCK2 | 10 | <10 | 20 | 10 | 80 | 80 | 320 | 320 | 80 | 320 | 10 | <10 | <10 | <10 | <10 | 10 | <10 | <10 | <10 | <10 | 320 | 320 |
| A/SWINE/MEXICO/AVX61/2013 | MDCK2 | 10 | 10 | 10 | 10 | 10 | 10 | 20 | 20 | 10 | 20 | 20 | 10 | 10 | 10 | 10 | <10 | <10 | <10 | 20 | 20 | 10 | 10 |
| A/SOLOMON_ISLANDS/3/2006 | E2 | <10 | <10 | * | <20 | 14 | 80 | <40 | * | 453 | 640 | * | * | <10 | <10 | * | * | <10 | <10 | * | * | 113 | 80 |
| A/SWINE/ALBERTA/SD0125/2015 | MDCK2 | * | * | <10 | <10 | <10 | * | <10 | <10 | <10 | <10 | * | * | 320 | 80 | * | * | * | * | * | * | <10 | <10 |
| A/SWINE/SASKATCHEWAN/SD0094/2015 | MDCK2 | * | * | <10 | 10 | 10 | * | <10 | <10 | <10 | 10 | * | * | 160 | 80 | * | * | * | * | * | * | <10 | <10 |
| A/SWINE/ALBERTA/SD0014/2013 | MDCK2 | * | * | <10 | <10 | <10 | * | <10 | <10 | <10 | <10 | * | * | 40 | 20 | * | * | * | * | * | * | <10 | <10 |
| A/SWINE/ALBERTA/SD0154/2016 | MDCK2 | * | * | <10 | <10 | <10 | * | <10 | <10 | <10 | <10 | * | * | 160 | 160 | * | * | * | * | * | * | <10 | <10 |
| A/SWINE/MANITOBA/D0333/2014 | MDCK2 | * | * | <10 | <10 | <10 | * | <10 | <10 | <10 | <10 | * | * | <10 | 10 | * | * | * | * | * | * | <10 | <10 |
| A/SWINE/MANITOBA/D0392/2014 | MDCK2 | * | * | <10 | <10 | <10 | * | <10 | <10 | <10 | <10 | * | * | <10 | <10 | * | * | * | * | * | * | <10 | <10 |
| A/SWINE/SASKATCHEWAN/SD0102/2015 | MDCK2 | * | * | <10 | <10 | <10 | * | <10 | <10 | <10 | <10 | * | * | <10 | <10 | * | * | * | * | * | * | <10 | <10 |
| A/SWINE/MANITOBA/SD0114/2015 | MDCK2 | * | * | <10 | <10 | <10 | * | <10 | <10 | <10 | <10 | * | * | <10 | <10 | * | * | * | * | * | * | <10 | <10 |
| A/SWINE/ALBERTA/SD0191/2016 | MDCK2 | * | * | <10 | <10 | <10 | * | <10 | <10 | <10 | <10 | * | * | <10 | <10 | * | * | * | * | * | * | 20 | 80 |
| A/SWINE/SASKATCHEWAN/SD0200/2016 | MDCK2 | * | * | 10 | <10 | <10 | * | 20 | 40 | <10 | <10 | * | * | <10 | <10 | * | * | * | * | * | * | 160 | 320 |
| A/SWINE/MANITOBA/D0348/2014 | MDCK2 | * | * | 10 | 10 | 10 | * | 80 | 160 | <10 | <10 | * | * | <10 | <10 | * | * | * | * | * | * | <10 | <10 |
| A/MINNESOTA/45/2016 | MDCK3 | * | * | <10 | <10 | <10 | * | <10 | <10 | <10 | <10 | * | * | <10 | <10 | * | * | * | * | * | * | <10 | <10 |
| A/SWINE/MINNESOTA/A01781045/2016 | MDCK2 | * | * | * | * | * | * | * | * | * | * | * | * | * | * | * | * | * | * | * | * | * | * |
| A/SINGAPORE/1986 | MDCK2 | 10 | 10 | 40 | 40 | 10 | <10 | 10 | 20 | 20 | 40 | * | * | * | * | * | * | <10 | 20 | * | * | 160 | 80 |
| A/TAIWAN/1/1986 | MDCK2 | 40 | 10 | 320 | 320 | <10 | <10 | <10 | <10 | <10 | <10 | * | * | * | * | * | * | <10 | <10 | * | * | <10 | 10 |
| A/TEXAS/36/1991 | MDCK2 | 10 | 10 | 320 | 320 | 20 | 20 | <10 | <10 | 10 | 10 | * | * | * | * | * | * | <10 | <10 | * | * | <10 | 20 |
| A/BEIJING/262/1995 | MDCK2 | <10 | <10 | 10 | 10 | 160 | 160 | 80 | 80 | 10 | 20 | * | * | * | * | * | * | <10 | <10 | * | * | 80 | 80 |
| A/MICHIGAN/2/2003 | MDCK2 | * | * | 20 | 20 | 28 | 40 | 80 | 80 | 40 | 80 | * | * | * | * | * | * | <10 | <10 | * | * | 160 | 226 |
| A/BRISBANE/59/2007 | MDCK3 | <10 | <10 | <10 | <10 | <10 | <10 | <10 | <10 | 80 | 80 | * | * | * | * | * | * | <10 | <10 | * | * | <10 | <10 |
| A/SWINE/NORTH_CAROLINA/A01730369/2016 | MDCK3 | <10 | <10 | <10 | <10 | <10 | <10 | <10 | <10 | <10 | <10 | * | * | * | * | * | * | 20 | <10 | * | * | 80 | 160 |
| A/SOLOMON_ISLANDS/3/2006 | E2/M1 | 20 | 10 | 40 | 20 | 10 | 40 | * | * | 320 | 320 | * | * | * | * | * | * | * | * | * | * | 40 | 40 |
| A/SWINE/MINNESOTA/A01134353/2011 | MDCK3 | 10 | <10 | <10 | <10 | <10 | <10 | * | * | <10 | <10 | * | * | * | * | * | * | * | * | * | * | 10 | 10 |
| A/SWINE/NEBRASKA/A01492366/2014 | MDCK3 | 40 | 40 | 40 | 20 | 20 | 40 | * | * | <10 | 20 | * | * | * | * | * | * | * | * | * | * | 160 | 80 |
| A/SWINE/ILLINOIS/A01644323/2018 | MDCK3 | 20 | 10 | 10 | 10 | 10 | 10 | * | * | <10 | 10 | * | * | * | * | * | * | * | * | * | * | 40 | 10 |
| A/SWINE/OKLAHOMA/A01409770/2014 | MDCK3 | 20 | 20 | 10 | 10 | 20 | <10 | * | * | 10 | 10 | * | * | * | * | * | * | * | * | * | * | 20 | 10 |
| A/SWINE/ILLINOIS/A02139356/2018 | MDCK3 | <10 | <10 | <10 | <10 | <10 | <10 | * | * | <10 | <10 | * | * | * | * | * | * | * | * | * | * | 10 | <10 |
| A/SWINE/MICHIGAN/A01104117/2018 | MDCK2 | 10 | 10 | 20 | 20 | 20 | 10 | * | * | 20 | 20 | * | * | * | * | * | * | * | * | * | * | 80 | 80 |

| Antigen names | Swine serum IDs | SD-P3 | SD-P4 | SD-P10 | SD-P11 | SD-P12 | SD-P15 | EXP480-P874 | EXP480-P875 | SD-P7 | SD-P8 | SD-P1 | SD-P2 | SD-P5 | SD-P6 | SD-P17 | SD-P18 | FLU9-P583 | FLU9-P585 | EXPXXX-P175 | EXPXXX-P176 | EXPXXX-P291 | EXPXXX-P292 | EXPXXX-P1 | EXPXXX-P2 |
|---|---|---|---|---|---|---|---|---|---|---|---|---|---|---|---|---|---|---|---|---|---|---|---|---|---|
| | Passage/Serum names | A/SWINE/ALBERTA/SD0125/2015 | A/SWINE/ALBERTA/SD0125/2015 | A/SWINE/ALBERTA/SD0014/2013 | A/SWINE/ALBERTA/SD0154/2016 | A/SWINE/ALBERTA/SD0154/2016 | A/SWINE/MANITOBA/D0333/2014 | A/SWINE/MINNESOTA/02053/2008 | A/SWINE/MINNESOTA/02053/2008 | A/SWINE/MANITOBA/D0392/2014 | A/SWINE/MANITOBA/D0392/2014 | A/SWINE/SASKATCHEWAN/SD0102/2015 | A/SWINE/SASKATCHEWAN/SD0102/2015 | A/SWINE/MANITOBA/SD0114/2015 | A/SWINE/MANITOBA/SD0114/2015 | A/SWINE/SASKATCHEWAN/SD0142/2016 | A/SWINE/SASKATCHEWAN/SD0142/2016 | A/SWINE/MINNESOTA/A01781045/2016 | A/SWINE/MINNESOTA/A01781045/2016 | A/SWINE/MINNESOTA/A01134353/2011 | A/SWINE/MINNESOTA/A01134353/2011 | A/SWINE/NEBRASKA/A01492366/2014 | A/SWINE/NEBRASKA/A01492366/2014 | A/SWINE/OKLAHOMA/A01409770/2014 | A/SWINE/OKLAHOMA/A01409770/2014 |
| A/SWINE/NORTH_CAROLINA/A02076926/2015 | MDCK2 | * | * | * | * | * | * | * | * | * | * | * | * | * | * | * | * | * | * | * | * | * | * | * | * |
| A/SWINE/NORTH_CAROLINA/A01841602/2015 | MDCK2 | * | * | * | * | * | * | * | * | * | * | * | * | * | * | * | * | * | * | * | * | * | * | * | * |
| A/SWINE/OHIO/A01847657/2015 | MDCK2 | * | * | * | * | * | * | * | * | * | * | * | * | * | * | * | * | * | * | * | * | * | * | * | * |
| A/OHIO/9/2015 | MDCK2 | * | * | * | * | * | * | 10 | <10 | * | * | * | * | * | * | * | * | * | * | * | * | * | * | * | * |
| A/SWINE/ILLINOIS/A01493472/2014 | MDCK2 | * | * | * | * | * | * | * | * | * | * | * | * | * | * | * | * | * | * | * | * | * | * | * | * |
| A/SWINE/MINNESOTA/A01567490/2014 | MDCK2 | * | * | * | * | * | * | * | * | * | * | * | * | * | * | * | * | * | * | * | * | * | * | * | * |
| A/CALIFORNIA/4/2009 | MDCK4 | * | * | * | * | * | * | 10 | <10 | * | * | * | * | * | * | * | * | * | * | * | * | * | * | * | * |
| A/MEXICO/4108/2009 | MDCK3 | * | * | * | * | * | * | * | * | * | * | * | * | * | * | * | * | * | * | * | * | * | * | * | * |
| A/BRAZIL/11/1978 | MDCK2 | <10 | <10 | <10 | <10 | <10 | <10 | <10 | <10 | <10 | <10 | <10 | <10 | <10 | <10 | <10 | <10 | * | * | <10 | <10 | <10 | <10 | 20 | 20 |
| A/NEW_CALEDONIA/20/1999 | E2 | <10 | <10 | <10 | <10 | <10 | <10 | <10 | <10 | <10 | <10 | <10 | <10 | <10 | <10 | <10 | <10 | * | * | * | * | * | * | * | * |
| A/MICHIGAN/45/2015 | MDCK3 | * | * | * | * | * | * | * | * | * | * | * | * | * | * | * | * | * | * | * | * | * | * | * | * |
| A/SWINE/NORTH_CAROLINA/A01797415/2015 | MDCK3 | * | * | * | * | * | * | * | * | * | * | * | * | * | * | * | * | * | * | * | * | * | * | * | * |
| A/SWINE/IOWA/A01731653/2016 | MDCK3 | * | * | * | * | * | * | * | * | * | * | * | * | * | * | * | * | * | * | * | * | * | * | * | * |
| A/SWINE/MEXICO/AVX31/2012 | MDCK2 | * | * | * | * | * | * | * | * | * | * | * | * | * | * | * | * | * | * | * | * | * | * | * | * |
| A/SWINE/MEXICO/AVX23/2012 | MDCK2 | * | * | * | * | * | * | * | * | * | * | * | * | * | * | * | * | * | * | * | * | * | * | * | * |
| A/SWINE/MEXICO/AVX44/2012 | MDCK2 | * | * | * | * | * | * | * | * | * | * | * | * | * | * | * | * | * | * | * | * | * | * | * | * |
| A/SWINE/MEXICO/AVX18/2012 | MDCK2 | * | * | * | * | * | * | * | * | * | * | * | * | * | * | * | * | * | * | * | * | * | * | * | * |
| A/SWINE/MEXICO/AVX61/2013 | MDCK2 | * | * | * | * | * | * | * | * | * | * | * | * | * | * | * | * | * | * | * | * | * | * | * | * |
| A/SOLOMON_ISLANDS/3/2006 | E2 | <10 | <10 | <10 | <10 | <10 | <10 | <10 | <10 | <10 | <10 | <10 | <10 | <10 | <10 | <10 | <10 | * | * | * | * | * | * | * | * |
| A/SWINE/ALBERTA/SD0125/2015 | MDCK2 | 640 | 160 | 10 | 80 | 160 | <10 | 40 | 80 | <10 | <10 | <10 | <10 | <10 | <10 | <10 | <10 | <10 | <10 | * | * | * | * | * | * |
| A/SWINE/SASKATCHEWAN/SD0094/2015 | MDCK2 | 80 | 20 | 20 | 40 | 160 | <10 | 160 | 160 | <10 | <10 | <10 | <10 | <10 | <10 | <10 | <10 | <10 | <10 | * | * | * | * | * | * |
| A/SWINE/ALBERTA/SD0014/2013 | MDCK2 | 40 | 10 | 320 | 20 | 80 | <10 | 20 | 40 | <10 | <10 | <10 | <10 | <10 | <10 | <10 | 10 | <10 | <10 | * | * | * | * | * | * |
| A/SWINE/ALBERTA/SD0154/2016 | MDCK2 | 80 | 20 | 10 | 160 | 640 | <10 | 40 | 40 | <10 | <10 | <10 | <10 | <10 | <10 | <10 | <10 | <10 | <10 | * | * | * | * | * | * |
| A/SWINE/MANITOBA/D0333/2014 | MDCK2 | <10 | <10 | <10 | <10 | <10 | 80 | <10 | <10 | <10 | 10 | <10 | <10 | <10 | <10 | 20 | <10 | <10 | <10 | * | * | * | * | * | * |
| A/SWINE/MANITOBA/D0392/2014 | MDCK2 | <10 | <10 | <10 | <10 | <10 | <10 | <10 | <10 | 80 | 40 | <10 | <10 | <10 | <10 | 20 | <10 | <10 | 10 | * | * | * | * | * | * |
| A/SWINE/SASKATCHEWAN/SD0102/2015 | MDCK2 | <10 | <10 | <10 | <10 | <10 | <10 | <10 | <10 | 160 | 80 | 320 | 320 | 40 | 20 | 320 | 160 | 20 | 80 | * | * | * | * | * | * |
| A/SWINE/MANITOBA/SD0114/2015 | MDCK2 | <10 | <10 | <10 | <10 | <10 | 10 | <10 | <10 | 320 | 160 | 160 | 160 | 640 | 160 | 80 | 320 | 40 | 80 | * | * | * | * | * | * |
| A/SWINE/ALBERTA/SD0191/2016 | MDCK2 | <10 | <10 | <10 | <10 | <10 | 20 | <10 | <10 | 320 | 640 | 640 | 640 | 80 | 40 | 320 | 320 | 40 | 160 | * | * | * | * | * | * |
| A/SWINE/SASKATCHEWAN/SD0200/2016 | MDCK2 | <10 | <10 | <10 | <10 | <10 | 10 | <10 | <10 | 640 | 640 | 320 | 640 | 80 | 160 | 1280 | 640 | 40 | 160 | * | * | * | * | * | * |
| A/SWINE/MANITOBA/D0348/2014 | MDCK2 | <10 | <10 | <10 | <10 | <10 | 10 | <10 | <10 | 320 | 320 | 320 | 320 | 160 | 80 | 640 | 160 | 80 | 160 | * | * | * | * | * | * |
| A/MINNESOTA/45/2016 | MDCK3 | <10 | <10 | <10 | <10 | <10 | <10 | <10 | <10 | 40 | 40 | 40 | 40 | 10 | 10 | 40 | 10 | 80 | 160 | * | * | * | * | * | * |
| A/SWINE/MINNESOTA/A01781045/2016 | MDCK2 | <10 | <10 | <10 | <10 | <10 | <10 | <10 | <10 | 40 | 40 | 20 | 20 | 40 | 20 | 10 | 40 | 160 | 320 | * | * | * | * | * | * |
| A/SINGAPORE/1986 | MDCK2 | * | * | * | * | * | * | 10 | 10 | * | * | * | * | * | * | * | * | * | * | * | * | * | * | * | * |
| A/TAIWAN/1/1986 | MDCK2 | * | * | * | * | * | * | <10 | <10 | * | * | * | * | * | * | * | * | * | * | * | * | * | * | * | * |
| A/TEXAS/36/1991 | MDCK2 | * | * | * | * | * | * | <10 | <10 | * | * | * | * | * | * | * | * | * | * | * | * | * | * | * | * |
| A/BEIJING/262/1995 | MDCK2 | * | * | * | * | * | * | <10 | <10 | * | * | * | * | * | * | * | * | * | * | * | * | * | * | * | * |
| A/MICHIGAN/2/2003 | MDCK2 | * | * | * | * | * | * | <10 | <10 | * | * | * | * | * | * | * | * | * | * | 40 | 40 | 80 | 160 | 20 | 10 |
| A/BRISBANE/59/2007 | MDCK3 | * | * | * | * | * | * | <10 | <10 | * | * | * | * | * | * | * | * | * | * | * | * | * | * | * | * |
| A/SWINE/NORTH_CAROLINA/A01730369/2016 | MDCK3 | * | * | * | * | * | * | 160 | 80 | * | * | * | * | * | * | * | * | * | * | * | * | * | * | * | * |
| A/SOLOMON_ISLANDS/3/2006 | E2/M1 | * | * | * | * | * | * | * | * | * | * | * | * | * | * | * | * | * | * | 20 | 40 | <10 | <10 | <10 | <10 |
| A/SWINE/MINNESOTA/A01134353/2011 | MDCK3 | * | * | * | * | * | * | * | * | * | * | * | * | * | * | * | * | * | * | 1280 | 640 | 40 | 40 | <10 | <10 |
| A/SWINE/NEBRASKA/A01492366/2014 | MDCK3 | * | * | * | * | * | * | * | * | * | * | * | * | * | * | * | * | * | * | 40 | 80 | 320 | 160 | 40 | 10 |
| A/SWINE/ILLINOIS/A01644323/2018 | MDCK3 | * | * | * | * | * | * | * | * | * | * | * | * | * | * | * | * | * | * | 1280 | 320 | 320 | 160 | 20 | <10 |
| A/SWINE/OKLAHOMA/A01409770/2014 | MDCK3 | * | * | * | * | * | * | * | * | * | * | * | * | * | * | * | * | * | * | 10 | 10 | 40 | 40 | 80 | 80 |
| A/SWINE/ILLINOIS/A02139356/2018 | MDCK3 | * | * | * | * | * | * | * | * | * | * | * | * | * | * | * | * | * | * | <10 | <10 | 10 | 20 | 20 | 20 |
| A/SWINE/MICHIGAN/A01104117/2018 | MDCK2 | * | * | * | * | * | * | * | * | * | * | * | * | * | * | * | * | * | * | 20 | 20 | 40 | 80 | 80 | 80 |

**Table S3.** Risk ranking scoring system used to prioritize representative swine H3N2 strains for the assessment of human immunity for pandemic potential.

| strain | global_lineage | blast_USA 2yr | blast_global 10yr | revblast_USA 2yr | revblast_global 10yr | **Sequence Factor** | A/BRAZIL/11/1978 | A/SINGAPORE/1986 | A/TAIWAN/1/1986 | A/TEXAS/36/1991 | A/BEIJING/262/1995 | A/NEW_CALEDONIA/20/1999 | A/MICHIGAN/2/2003 | A/SOLOMON_ISLANDS/3/2006 | A/BRISBANE/59/2007 | A/CALIFORNIA/4/2009 | A/MICHIGAN/45/2015 | all_vaccine | stdev | >3 | **Antigenc Factor** | **Summed Factor** |
|---|---|---|---|---|---|---|---|---|---|---|---|---|---|---|---|---|---|---|---|---|---|---|
| **A/SWINE/SASKATCHEWAN/SD0094/2015** | **1A.1.1-1** | **0.00** | **0.18** | **0.00** | **23.67** | **40.40** | **5.77** | **3.65** | **5.82** | **5.62** | **6.40** | **5.69** | **5.09** | **6.15** | **7.81** | **3.54** | **2.56** | **5.28** | **1.49** | **10** | **14.48** | **54.89** |
| A/SWINE/ALBERTA/SD0154/2016 | 1A.1.1-1 | 0.00 | 0.06 | 0.00 | 0.06 | 0.20 | 6.96 | 4.91 | 6.62 | 6.41 | 7.56 | 6.25 | 6.27 | 6.69 | 7.79 | 2.12 | 1.85 | 5.77 | 2.01 | 9 | 15.04 | 15.25 |
| A/SWINE/ALBERTA/SD0125/2015 | 1A.1.1-1 | 0.00 | 0.06 | 0.00 | 9.45 | 16.11 | 6.03 | 5.29 | 6.05 | 6.09 | 7.99 | 7.37 | 6.77 | 7.34 | 8.65 | 3.12 | 1.75 | 6.04 | 2.05 | 10 | 16.16 | 32.27 |
| A/SWINE/ALBERTA/SD0014/2013 | 1A.1.1-1 | 0.00 | 0.06 | 0.00 | 0.06 | 0.20 | 6.91 | 5.54 | 8.09 | 8.08 | 8.23 | 7.83 | 6.93 | 8.79 | 10.56 | 4.35 | 2.47 | 7.07 | 2.24 | 10 | 16.71 | 16.92 |
| A/SWINE/MANITOBA/D0333/2014 | 1A.1.1-3 | 0.00 | 0.06 | 0.00 | 0.18 | 0.41 | 9.33 | 5.22 | 9.49 | 8.96 | 6.89 | 5.00 | 5.72 | 7.43 | 8.87 | 5.87 | 5.92 | 7.15 | 1.74 | 11 | 16.21 | 16.62 |
| A/SWINE/SASKATCHEWAN/SD0200/2016 | 1A.1.1-3-del | 0.00 | 0.12 | 0.00 | 0.18 | 0.51 | 6.17 | 3.33 | 7.13 | 6.75 | 2.97 | 5.34 | 2.77 | 6.42 | 9.32 | 9.62 | 8.65 | 6.22 | 2.44 | 9 | 16.33 | 16.84 |
| A/SWINE/MANITOBA/D0348/2014 | 1A.1.1-3-del | 0.00 | 0.24 | 0.22 | 1.65 | 3.50 | 5.82 | 3.38 | 6.90 | 6.56 | 2.96 | 5.64 | 2.86 | 6.49 | 9.45 | 9.80 | 8.75 | 6.24 | 2.47 | 9 | 16.40 | 19.90 |
| A/SWINE/MANITOBA/SD0114/2015 | 1A.1.1-3-del | 0.00 | 0.41 | 3.51 | 1.06 | 7.29 | 5.70 | 4.29 | 7.65 | 7.52 | 4.83 | 7.11 | 4.42 | 8.05 | 10.97 | 9.75 | 8.34 | 7.15 | 2.17 | 11 | 17.51 | 24.80 |
| A/SWINE/ALBERTA/SD0191/2016 | 1A.1.1-3-del | 0.00 | 0.12 | 0.15 | 0.41 | 1.10 | 6.35 | 3.90 | 7.61 | 7.28 | 3.62 | 6.01 | 3.46 | 7.11 | 10.02 | 10.00 | 8.92 | 6.75 | 2.38 | 11 | 18.13 | 19.24 |
| A/SWINE/SASKATCHEWAN/SD0102/2015 | 1A.1.1-3-del | 0.00 | 0.12 | 0.15 | 0.83 | 1.81 | 7.38 | 5.82 | 8.91 | 8.66 | 4.88 | 7.73 | 5.18 | 8.64 | 11.61 | 12.07 | 10.89 | 8.34 | 2.47 | 11 | 18.42 | 20.24 |
| **A/SWINE/MINNESOTA/A01781045/2016** | **1A.1.1-3-del** | **0.00** | **0.18** | **2.71** | **1.36** | **6.31** | **3.25** | **4.13** | **5.14** | **5.21** | **4.10** | **7.39** | **4.19** | **7.01** | **10.02** | **10.30** | **8.90** | **6.33** | **2.53** | **11** | **18.59** | **24.91** |
| A/MINNESOTA/45/2016 | 1A.1.1-3-del | 1.10 | 0.00 | 5.34 | 0.47 | 9.60 | 4.77 | 5.29 | 6.18 | 6.15 | 4.25 | 7.95 | 4.84 | 7.53 | 10.56 | 11.86 | 10.59 | 7.27 | 2.67 | 11 | 19.00 | 28.60 |
| A/SWINE/MANITOBA/D0392/2015 | 1A.1.1-3-del | 0.00 | 0.12 | 0.00 | 0.24 | 0.61 | 8.06 | 7.03 | 8.07 | 7.69 | 4.53 | 7.95 | 5.81 | 7.50 | 10.20 | 13.91 | 13.14 | 8.54 | 2.85 | 11 | 19.54 | 20.15 |
| **A/SWINE/MEXICO/AVX23/2012** | **1A.3.1** | **0.00** | **0.24** | **2.71** | **7.14** | **16.21** | **7.35** | **6.23** | **6.25** | **6.15** | **8.64** | **7.34** | **7.51** | **6.93** | **7.42** | **2.79** | **3.20** | **6.35** | **1.81** | **10** | **15.43** | **31.64** |
| A/SWINE/MEXICO/AVX31/2012 | 1A.3.3.2 | 0.00 | 0.06 | 0.00 | 3.25 | 5.61 | 7.75 | 5.66 | 7.05 | 6.80 | 8.20 | 6.51 | 6.93 | 6.87 | 7.55 | 1.45 | 2.27 | 6.10 | 2.20 | 9 | 15.61 | 21.21 |
| A/SWINE/MEXICO/AVX44/2012 | 1A.3.3.2 | 0.00 | 0.12 | 0.00 | 0.18 | 0.51 | 7.73 | 4.59 | 7.95 | 7.62 | 7.08 | 5.62 | 5.72 | 7.12 | 8.57 | 3.58 | 3.21 | 6.25 | 1.83 | 11 | 16.50 | 17.00 |
| A/SWINE/ILLINOIS/A01493472/2014 | 1A.3.3.2 | 0.00 | 0.35 | 6.88 | 30.70 | 62.00 | 9.14 | 7.22 | 8.42 | 8.21 | 9.73 | 7.77 | 8.46 | 8.19 | 8.44 | 0.71 | 2.72 | 7.18 | 2.82 | 9 | 17.45 | 79.45 |
| A/SWINE/NORTH_CAROLINA/A01797415/2015 | 1A.3.3.3 | 0.00 | 0.00 | 0.22 | 0.30 | 0.81 | 7.33 | 5.00 | 5.59 | 5.14 | 7.02 | 5.24 | 5.97 | 4.83 | 5.23 | 3.74 | 4.53 | 5.42 | 1.04 | 11 | 14.12 | 14.93 |
| **A/SWINE/IOWA/A01731653/2016** | **1A.3.3.3** | **1.68** | **0.00** | **36.21** | **0.12** | **52.00** | **6.82** | **5.54** | **5.05** | **4.81** | **7.63** | **6.43** | **6.64** | **5.53** | **6.00** | **4.09** | **4.49** | **5.73** | **1.09** | **11** | **14.26** | **66.26** |
| A/SWINE/NORTH_CAROLINA/A01730369/2016 | 1A.3.3.3 | 0.00 | 0.00 | 0.00 | 0.00 | 0.00 | 6.19 | 2.77 | 6.53 | 6.18 | 5.33 | 4.69 | 3.98 | 5.95 | 7.96 | 4.86 | 4.08 | 5.32 | 1.44 | 10 | 14.32 | 14.32 |
| A/SWINE/MINNESOTA/A01567490/2014 | 1A.3.3.3 | 0.07 | 0.00 | 2.56 | 1.53 | 6.19 | 8.40 | 6.39 | 6.98 | 6.66 | 8.63 | 6.65 | 7.49 | 6.53 | 6.55 | 2.51 | 3.88 | 6.42 | 1.79 | 10 | 15.38 | 21.56 |
| A/OHIO/09/2015 | 1A.3.3.3 | 0.00 | 0.00 | 0.00 | 0.06 | 0.10 | 9.26 | 5.38 | 8.00 | 7.35 | 7.09 | 3.99 | 5.97 | 5.40 | 5.59 | 4.14 | 5.43 | 6.14 | 1.62 | 11 | 15.85 | 15.95 |
| A/SWINE/NORTH_CAROLINA/A02076926/2015 | 1A.3.3.3 | 0.07 | 0.00 | 1.46 | 0.06 | 2.19 | 10.44 | 7.07 | 8.92 | 8.34 | 8.86 | 5.74 | 7.76 | 6.64 | 6.02 | 3.75 | 5.68 | 7.20 | 1.89 | 11 | 16.68 | 18.88 |
| A/SWINE/NORTH_CAROLINA/A01841602/2015 | 1A.3.3.3 | 0.00 | 0.00 | 0.80 | 0.00 | 1.09 | 10.84 | 7.00 | 8.98 | 8.23 | 8.18 | 4.67 | 7.30 | 5.68 | 4.62 | 5.62 | 7.29 | 7.13 | 1.91 | 11 | 16.72 | 17.81 |
| A/SWINE/OHIO/A01847657/2015 | 1A.3.3.3 | 0.00 | 0.00 | 0.00 | 0.06 | 0.10 | 10.82 | 6.62 | 9.30 | 8.53 | 7.74 | 4.09 | 6.83 | 5.80 | 5.31 | 5.80 | 7.28 | 7.10 | 1.93 | 11 | 16.78 | 16.88 |
| A/SWINE/MICHIGAN/A01104117/2018 | 1B.2.1 | 1.83 | 0.00 | 5.41 | 0.00 | 9.90 | 4.32 | 2.01 | 4.75 | 4.38 | 1.94 | 4.91 | 1.71 | 4.91 | 7.91 | 8.84 | 7.90 | 4.87 | 2.48 | 8 | 15.44 | 25.34 |
| A/SWINE/OKLAHOMA/A01409770/2014 | 1B.2.1 | 0.15 | 0.00 | 2.56 | 2.01 | 7.11 | 3.60 | 2.89 | 4.84 | 4.70 | 3.00 | 6.10 | 2.90 | 5.94 | 8.96 | 9.37 | 8.16 | 5.50 | 2.43 | 9 | 16.29 | 23.40 |
| **A/SWINE/ILLINOIS/A02139356/2018** | **1B.2.1** | **0.37** | **0.00** | **8.34** | **0.00** | **11.91** | **4.84** | **4.51** | **6.47** | **6.37** | **3.85** | **7.29** | **4.15** | **7.36** | **10.43** | **11.02** | **9.72** | **6.91** | **2.55** | **11** | **18.65** | **30.55** |
| A/SWINE/ILLINOIS/A01644323/2018 | 1B.2.2.1 | 0.07 | 0.00 | 6.36 | 1.06 | 10.59 | 5.37 | 1.68 | 5.81 | 5.45 | 4.24 | 4.39 | 2.92 | 5.40 | 7.79 | 5.90 | 5.01 | 4.91 | 1.61 | 9 | 13.83 | 24.42 |
| **A/SWINE/NEBRASKA/A01492366/2014** | **1B.2.2.1** | **0.07** | **0.00** | **8.71** | **0.65** | **13.10** | **4.42** | **0.53** | **4.62** | **4.20** | **2.73** | **4.20** | **1.61** | **4.51** | **7.29** | **7.24** | **6.38** | **4.34** | **2.15** | **8** | **14.44** | **27.54** |
| A/SWINE/MINNESOTA/A01134353/2011 | 1B.2.2.1 | 0.00 | 0.00 | 5.71 | 0.00 | 7.81 | 6.64 | 3.28 | 6.85 | 6.49 | 5.82 | 4.81 | 4.46 | 6.13 | 7.93 | 4.32 | 3.70 | 5.49 | 1.47 | 11 | 15.41 | 23.22 |
| **A/SWINE/MEXICO/AVX18/2012** | **Other_Human_1B.2** | **0.00** | **0.12** | **0.00** | **12.63** | **21.60** | **6.87** | **2.65** | **5.71** | **4.88** | **1.96** | **1.89** | **1.60** | **2.65** | **5.29** | **8.23** | **8.15** | **4.53** | **2.52** | **6** | **13.56** | **35.16** |
| A/SWINE/MEXICO/AVX61/2013 | Other_Human_1B.2 | 0.00 | 0.06 | 0.00 | 0.59 | 1.10 | 5.01 | 1.21 | 5.53 | 5.16 | 3.65 | 4.36 | 2.39 | 5.23 | 7.80 | 6.55 | 5.62 | 4.77 | 1.84 | 9 | 14.52 | 15.62 |

**Table S4.** Hemagglutination inhibition titres of post-infection convalescent (post_exp) and post-vaccination (post_vac) human sera against representative North American swine H1strains.

| study_id | cohort | hospital | age | yearofbirth | periodofbirth | gender | pre_vaccine_titre | post_vaccine_titre | fold_change | BE95.huvac | MI15.huvac | 1A.1.1.1 SK15 | 1A.1.1.3 MN16 | 1A.3.3.3 IA16 | 1A.3.1 MX12 | OtHu1B.2 MX12 | 1B.2.2.1 NE14 | 1B.2.1 IL18 |
|---|---|---|---|---|---|---|---|---|---|---|---|---|---|---|---|---|---|---|
| 01-11-A-0077 | post_exp | JHHS | 71 | 1946 | 1946 to 1976 | n/a | n/a | n/a | n/a | 10 | 320 | 80 | 5 | 80 | 20 | 10 | 20 | 5 |
| 02-17-Pro-0089 | post_vac | JHHS | 68 | 1949 | 1946 to 1976 | Female | 80 | 320 | 4 | 5 | 20 | 20 | 5 | 5 | 5 | 10 | 20 | 5 |
| 02-17-Pro-0027 | post_vac | JHHS | 66 | 1951 | 1946 to 1976 | Male | 320 | 640 | 2 | 40 | 160 | 10 | 5 | 40 | 40 | 5 | 160 | 40 |
| 01-21-A-0046 | post_exp | CGMH | 65 | 1952 | 1946 to 1976 | n/a | n/a | n/a | n/a | 10 | 40 | 20 | 5 | 20 | 40 | 10 | 80 | 10 |
| 02-17-Pro-0037 | post_vac | JHHS | 61 | 1956 | 1946 to 1976 | Male | 1280 | 2560 | 2 | 2560 | 640 | 640 | 5 | 160 | 80 | 2560 | 320 | 20 |
| 02-17-Pro-0111 | post_vac | JHHS | 59 | 1958 | 1946 to 1976 | Female | 2560 | 5120 | 2 | 5 | 160 | 20 | 5 | 40 | 10 | 10 | 10 | 5 |
| 02-17-Pro-0121 | post_vac | JHHS | 59 | 1958 | 1946 to 1976 | Male | 640 | 1280 | 2 | 640 | 40 | 20 | 10 | 20 | 10 | 1280 | 320 | 10 |
| 02-17-Pro-0124 | post_vac | JHHS | 58 | 1959 | 1946 to 1976 | Female | 10240 | 10240 | 1 | 10 | 160 | 80 | 40 | 40 | 20 | 40 | 160 | 80 |
| 02-17-Pro-0051 | post_vac | JHHS | 56 | 1961 | 1946 to 1976 | Female | 80 | 640 | 8 | 40 | 160 | 80 | 10 | 160 | 80 | 160 | 80 | 5 |
| 02-17-Pro-0012 | post_vac | JHHS | 53 | 1964 | 1946 to 1976 | Female | 640 | 1280 | 2 | 80 | 320 | 10 | 5 | 80 | 10 | 80 | 40 | 5 |
| 02-17-Pro-0016 | post_vac | JHHS | 51 | 1966 | 1946 to 1976 | Male | 320 | 640 | 2 | 80 | 640 | 80 | 5 | 640 | 80 | 80 | 80 | 5 |
| 02-17-Pro-0080 | post_vac | JHHS | 51 | 1966 | 1946 to 1976 | Male | 1280 | 1280 | 1 | 20 | 40 | 40 | 5 | 80 | 20 | 20 | 10 | 5 |
| 02-17-Pro-0100 | post_vac | JHHS | 49 | 1968 | 1946 to 1976 | Male | 40 | 320 | 8 | 10 | 10 | 80 | 5 | 80 | 160 | 80 | 20 | 5 |
| 02-17-Pro-0109 | post_vac | JHHS | 49 | 1968 | 1946 to 1976 | Male | 2560 | 2560 | 1 | 20 | 20 | 80 | 5 | 20 | 10 | 20 | 20 | 10 |
| 02-17-Pro-0103 | post_vac | JHHS | 48 | 1969 | 1946 to 1976 | Male | 1280 | 1280 | 1 | 10 | 40 | 320 | 5 | 160 | 320 | 20 | 40 | 10 |
| 02-17-Pro-0022 | post_vac | JHHS | 47 | 1970 | 1946 to 1976 | Male | 320 | 1280 | 4 | 40 | 160 | 20 | 5 | 40 | 10 | 20 | 10 | 5 |
| 01-21-A-0039 | post_exp | CGMH | 47 | 1970 | 1946 to 1976 | n/a | n/a | n/a | n/a | 640 | 320 | 640 | 5 | 320 | 80 | 640 | 80 | 5 |
| 02-17-Pro-0052 | post_vac | JHHS | 44 | 1973 | 1946 to 1976 | Female | 2560 | 5120 | 2 | 640 | 160 | 80 | 5 | 80 | 20 | 1280 | 320 | 5 |
| 02-17-Pro-0116 | post_vac | JHHS | 44 | 1973 | 1946 to 1976 | Male | 160 | 640 | 4 | 40 | 20 | 10 | 5 | 10 | 5 | 40 | 40 | 5 |
| 02-17-Pro-0004 | post_vac | JHHS | 42 | 1975 | 1946 to 1976 | Female | 5120 | 10240 | 2 | 160 | 640 | 320 | 5 | 160 | 160 | 20 | 20 | 5 |
| 02-17-Pro-0049 | post_vac | JHHS | 42 | 1975 | 1946 to 1976 | Female | 1280 | 2560 | 2 | 80 | 20 | 20 | 5 | 20 | 5 | 160 | 80 | 10 |
| 02-17-Pro-0104 | post_vac | JHHS | 41 | 1976 | 1946 to 1976 | Female | 80 | 160 | 2 | 80 | 20 | 160 | 5 | 40 | 20 | 160 | 40 | 5 |
| 02-17-Pro-0093 | post_vac | JHHS | 40 | 1977 | 1977 to 1988 | Female | 10240 | 10240 | 1 | 5 | 320 | 40 | 10 | 80 | 20 | 10 | 20 | 10 |
| 01-11-A-0065 | post_exp | JHHS | 40 | 1977 | 1977 to 1988 | n/a | n/a | n/a | n/a | 640 | 640 | 80 | 5 | 1280 | 640 | 80 | 1280 | 2560 |
| 02-17-Pro-0015 | post_vac | JHHS | 39 | 1978 | 1977 to 1988 | Female | 320 | 320 | 1 | 160 | 80 | 40 | 10 | 40 | 40 | 160 | 80 | 20 |
| 02-17-Pro-0114 | post_vac | JHHS | 39 | 1978 | 1977 to 1988 | Male | 160 | 320 | 2 | 80 | 160 | 160 | 20 | 160 | 160 | 80 | 10 | 5 |
| 01-11-A-0080 | post_exp | JHHS | 39 | 1978 | 1977 to 1988 | n/a | n/a | n/a | n/a | 160 | 80 | 40 | 5 | 80 | 160 | 160 | 80 | 80 |
| 02-17-Pro-0091 | post_vac | JHHS | 38 | 1979 | 1977 to 1988 | Female | 160 | 640 | 4 | 10 | 2560 | 160 | 5 | 320 | 80 | 40 | 40 | 5 |
| 02-17-Pro-0101 | post_vac | JHHS | 38 | 1979 | 1977 to 1988 | Male | 5120 | 10240 | 2 | 5 | 40 | 10 | 5 | 20 | 10 | 10 | 10 | 5 |
| 01-22-A-0070 | post_exp | CGMH | 38 | 1979 | 1977 to 1988 | n/a | n/a | n/a | n/a | 10 | 40 | 20 | 5 | 10 | 5 | 10 | 10 | 5 |
| 01-11-A-0116 | post_exp | JHHS | 36 | 1981 | 1977 to 1988 | n/a | n/a | n/a | n/a | 320 | 2560 | 1280 | 40 | 2560 | 2560 | 320 | 320 | 320 |
| 02-17-Pro-0070 | post_vac | JHHS | 34 | 1983 | 1977 to 1988 | Female | 2560 | 2560 | 1 | 10 | 320 | 640 | 5 | 160 | 160 | 10 | 5 | 5 |
| 02-17-Pro-0115 | post_vac | JHHS | 34 | 1983 | 1977 to 1988 | Male | 320 | 640 | 2 | 40 | 80 | 320 | 5 | 160 | 160 | 40 | 20 | 40 |
| 01-11-A-0121 | post_exp | JHHS | 34 | 1983 | 1977 to 1988 | n/a | n/a | n/a | n/a | 160 | 1280 | 1280 | 5 | 160 | 320 | 320 | 20 | 5 |
| 01-12-A-0053 | post_exp | JHHS | 34 | 1983 | 1977 to 1988 | n/a | n/a | n/a | n/a | 10 | 40 | 40 | 5 | 80 | 20 | 10 | 10 | 5 |
| 02-17-Pro-0013 | post_vac | JHHS | 33 | 1984 | 1977 to 1988 | Female | 640 | 640 | 1 | 20 | 320 | 40 | 5 | 160 | 40 | 160 | 40 | 10 |
| 01-12-A-0136 | post_exp | JHHS | 33 | 1984 | 1977 to 1988 | n/a | n/a | n/a | n/a | 80 | 160 | 160 | 10 | 160 | 80 | 160 | 20 | 5 |
| 01-21-A-0029 | post_exp | CGMH | 32 | 1985 | 1977 to 1988 | n/a | n/a | n/a | n/a | 160 | 320 | 160 | 5 | 160 | 160 | 320 | 160 | 5 |
| 02-17-Pro-0023 | post_vac | JHHS | 31 | 1986 | 1977 to 1988 | Male | 160 | 320 | 2 | 80 | 80 | 40 | 5 | 40 | 40 | 80 | 20 | 20 |
| 02-17-Pro-0069 | post_vac | JHHS | 31 | 1986 | 1977 to 1988 | Female | 1280 | 2560 | 2 | 20 | 320 | 160 | 5 | 160 | 160 | 20 | 10 | 5 |
| 02-17-Pro-0020 | post_vac | JHHS | 30 | 1987 | 1977 to 1988 | Male | 80 | 160 | 2 | 640 | 640 | 160 | 10 | 320 | 80 | 1280 | 640 | 40 |
| 01-21-A-0047 | post_exp | CGMH | 30 | 1987 | 1977 to 1988 | n/a | n/a | n/a | n/a | 320 | 320 | 160 | 10 | 160 | 80 | 320 | 320 | 10 |
| 02-17-Pro-0019 | post_vac | JHHS | 29 | 1988 | 1977 to 1988 | Female | 2560 | 2560 | 1 | 320 | 320 | 80 | 5 | 2560 | 2560 | 320 | 640 | 80 |
| 02-17-Pro-0083 | post_vac | JHHS | 29 | 1988 | 1977 to 1988 | Female | 2560 | 2560 | 1 | 40 | 320 | 640 | 5 | 160 | 80 | 40 | 20 | 5 |
| 01-21-A-0032 | post_exp | CGMH | 29 | 1988 | 1977 to 1988 | n/a | n/a | n/a | n/a | 160 | 20 | 40 | 5 | 10 | 10 | 80 | 40 | 5 |
| 01-22-A-0025 | post_exp | CGMH | 28 | 1989 | 1989 to 1996 | n/a | n/a | n/a | n/a | 40 | 160 | 80 | 5 | 20 | 40 | 40 | 10 | 5 |
| 02-17-Pro-0029 | post_vac | JHHS | 27 | 1990 | 1989 to 1996 | Male | 640 | 2560 | 4 | 80 | 80 | 40 | 5 | 20 | 20 | 320 | 40 | 5 |
| 01-11-A-0102 | post_exp | JHHS | 26 | 1991 | 1989 to 1996 | n/a | n/a | n/a | n/a | 320 | 320 | 640 | 40 | 320 | 320 | 320 | 320 | 40 |
| 02-17-Pro-0063 | post_vac | JHHS | 25 | 1992 | 1989 to 1996 | Female | 2560 | 5120 | 2 | 640 | 2560 | 160 | 10 | 1280 | 160 | 2560 | 1280 | 20 |
| 02-17-Pro-0095 | post_vac | JHHS | 25 | 1992 | 1989 to 1996 | Male | 320 | 1280 | 4 | 40 | 160 | 160 | 5 | 80 | 80 | 80 | 40 | 5 |
| 01-21-A-0077 | post_exp | CGMH | 25 | 1992 | 1989 to 1996 | n/a | n/a | n/a | n/a | 640 | 1280 | 640 | 5 | 320 | 640 | 640 | 320 | 20 |
| 01-11-A-0160 | post_exp | JHHS | 24 | 1993 | 1989 to 1996 | n/a | n/a | n/a | n/a | 80 | 5 | 5 | 5 | 5 | 5 | 160 | 160 | 5 |
| 02-17-Pro-0011 | post_vac | JHHS | 23 | 1994 | 1989 to 1996 | Female | 1280 | 2560 | 2 | 80 | 640 | 40 | 10 | 320 | 160 | 160 | 40 | 5 |
| 02-17-Pro-0043 | post_vac | JHHS | 23 | 1994 | 1989 to 1996 | Male | 10240 | 10240 | 1 | 40 | 320 | 5 | 5 | 10 | 5 | 40 | 5 | 5 |
| 02-17-Pro-0073 | post_vac | JHHS | 23 | 1994 | 1989 to 1996 | Male | 2560 | 2560 | 1 | 20 | 160 | 20 | 5 | 40 | 20 | 80 | 20 | 5 |
| 02-17-Pro-0096 | post_vac | JHHS | 23 | 1994 | 1989 to 1996 | Female | 10240 | 20480 | 2 | 20 | 80 | 40 | 5 | 80 | 80 | 20 | 40 | 20 |
| 02-17-Pro-0047 | post_vac | JHHS | 22 | 1995 | 1989 to 1996 | Male | 1280 | 1280 | 1 | 160 | 320 | 160 | 5 | 320 | 80 | 320 | 40 | 5 |
| 01-21-A-0083 | post_exp | CGMH | 22 | 1995 | 1989 to 1996 | n/a | n/a | n/a | n/a | 320 | 320 | 80 | 5 | 160 | 40 | 640 | 160 | 10 |
| 01-11-A-0026 | post_exp | JHHS | 21 | 1996 | 1989 to 1996 | n/a | n/a | n/a | n/a | 80 | 640 | 80 | 20 | 640 | 160 | 320 | 80 | 20 |
| 01-21-A-0060 | post_exp | CGMH | n/a | n/a | n/a | n/a | n/a | n/a | n/a | 5 | 640 | 20 | 5 | 80 | 10 | 5 | 20 | 5 |
